# Network rewiring: physiological consequences of reciprocally exchanging the physical locations and growth-phase-dependent expression patterns of the *Salmonella fis* and *dps* genes

**DOI:** 10.1101/2020.07.08.193441

**Authors:** Marina M Bogue, Aalap Mogre, Michael C Beckett, Nicholas R Thomson, Charles J Dorman

## Abstract

The Fis nucleoid-associated protein controls the expression of a large and diverse regulon of genes in Gram-negative bacteria. Fis production is normally maximal in bacteria during the early exponential phase of batch culture growth, becoming almost undetectable by the onset of stationary phase. We tested the effect on the Fis regulatory network in *Salmonella* of moving the complete *fis* gene from its usual location near the origin of chromosomal replication to the position normally occupied by the *dps* gene in the Right macrodomain of the chromosome, and *vice versa*, creating the strain GX. In a parallel experiment, we tested the effect of rewiring the Fis regulatory network by placing the *fis* open reading frame under the control of the stationary-phase-activated *dps* promoter at the *dps* genetic location within Ter, and *vice versa*, creating the strain OX. ChIP-seq was used to measure global Fis protein binding and gene expression patterns. Strain GX showed few changes when compared with the wild type, although we did detect increased Fis binding at Ter, accompanied by reduced binding at Ori. Strain OX displayed a more pronounced version of this distorted Fis protein-binding pattern together with numerous alterations in the expression of genes in the Fis regulon. OX, but not GX, had a reduced ability to infect cultured mammalian cells. These findings illustrate the inherent robustness of the Fis regulatory network to rewiring based on gene repositioning alone and emphasise the importance of *fis* expression signals in phenotypic determination.

**IMPORTANCE:** We assessed the impacts on *Salmonella* physiology of reciprocally translocating the genes encoding the Fis and Dps nucleoid-associated proteins (NAPs), and of inverting their growth phase production patterns such that Fis is produced in stationary phase (like Dps) and Dps is produced in exponential phase (like Fis). Changes to peak binding of Fis were detected by ChIP-seq on the chromosome, as were widespread impacts on the transcriptome, especially when Fis production mimicked Dps. Virulence gene expression and the expression of a virulence phenotype were altered. Overall, these radical changes to NAP gene expression were well tolerated, revealing the robust and well-buffered nature of global gene regulation networks in the bacterium.

## INTRODUCTION

Trans-acting regulatory proteins play a prominent role in controlling the expression of bacterial genes at the level of transcription. Whether acting positively or negatively, these proteins typically influence their target genes by binding at, or close to, transcriptional promoters (1). Transcription factors differ in the number of genes that are under their control (2) and a majority of these proteins exist with moderate copy numbers of between one and 100 monomers (3). Nucleoid-associated proteins (NAPs) play a role in the architecture of the genome and many contribute to the regulation of transcription (4). NAPs share many of the features of transcription factors but many are present in thousands of copies. The distinction between ‘transcription factor’ and ‘NAP’ is quite blurred and these are operational descriptions for proteins that lie along a continuum extending from factors with pervasive influences on gene expression to those that have just a few gene targets (5).

Transcription factors must make the journey from the gene that encodes them to the target promoter. This process seems to involve a combination of sliding along DNA and movements between DNA segments in which the interactions with DNA consist of non-specific and specific binding events (6, 7). In the case of a low-copy number transcription factor such as the Lac repressor protein, LacI (40 monomers per cell), the addition of an inducer (IPTG) reduces specific, but not the non-specific binding: the protein explores the nucleoid as it searches for its specific binding sites (6). Many NAPs rely on indirect readout to identify their DNA targets, making their binding site selections based on DNA conformation rather than on base sequence alone (5). This, combined with their high copy numbers, might allow NAPs to spread rapidly through the nucleoid to locate their DNA targets. Alternatively, their weak requirement for specific base sequences at their DNA targets might increase their dwell time at the many non-specific sites they encounter as they migrate through the nucleoid.

The significance of the genomic positions of genes encoding transcription factors or NAPs has been investigated, principally in the model bacterium *Escherichia coli*. There, a correlation is seen between gene position along the replichore extending from the origin of chromosome replication, *oriC*, to the terminus, and the period in the growth cycle when the gene product is most required. Gene position also correlates with gene copy number in fast-growing bacteria, when replication recommences before cell division is complete. This correlation is conserved across the gamma-Proteobacteria (8) suggesting that it is biologically meaningful and may have consequences for the operation of bacterial gene control networks. This hypothesis is supported by the observations that gene relocations that occur naturally, e.g. through inversions of chromosomal segments, tend to preserve the distance of the affected gene(s) from *oriC* (9–11). It suggests that moving regulatory genes, including genes that encode NAPs, to novel chromosomal locations could result in changes to bacterial physiology. Consistent with this proposal, repositioning the gene encoding the Factor for Inversion Stimulation (FIS) NAP in *E. coli* alters the cell’s capacity to manage its global DNA topology (12).

Fis, the Factor for Inversion Stimulation, is a prominent member of the family of NAPs in Gram-negative bacteria (13, 14) that alters the transcription of hundreds of genes, directly or indirectly, positively or negatively (15–19), and contributes architecturally to site-specific recombination systems (20–22), chromosome replication initiation (23–26), transposon activity (27, 28) and bacteriophage life cycles (22, 27, 29, 30). Fis is not essential, despite its pervasive influence on cell biology, but it enhances the fitness of a wild type bacterium when competing with an otherwise isogenic *fis* knockout mutant (31). Among the genes that are regulated by Fis are those encoding components of the translational apparatus of the bacterium, its chemotaxis and motility functions, many metabolic pathway proteins and (in the case of pathogens) numerous virulence genes (17, 19, 32–34). The association of high Fis concentrations with the early exponential phase of growth is thought to be indicative of a role for Fis in signalling growth-cycle-related information to the global gene expression programme of the cell (31, 35, 36).

The Fis protein influences the topology of DNA both directly, through DNA binding (31) and indirectly, through its influence on the expression of the genes encoding DNA gyrase (*gyrA* and *gyrB*) and DNA topoisomerase I (*topA*) (37–39). The *fis* gene is part of the bicistronic *dusB-fis* operon (40, 41) whose stringently controlled promoter is stimulated by DNA negative supercoiling, creating a regulatory connection between the global supercoiling level in bacterial DNA, bacterial physiology and the initiation of *fis* transcription (42). The single *dusB-fis* promoter is auto-repressed by the Fis protein (43) with translation of the Fis protein relying on mRNA secondary structure and nucleotide sequence motifs in the upstream message (44).

The production of high levels of Fis protein is associated with the early exponential phase of bacterial growth with Fis being present at very low levels in the stationary phase (35, 37, 45, 46). This pattern is sensitive to aeration: cultures of *Salmonella enterica* serovar Typhimurium exhibit sustained, low-level, production of Fis into the stationary phase of growth in the absence of aeration (47, 48).

We explored the subtleties of Fis biology by testing the significance of the geographical location of the *fis* gene in the Non-structured Left (NS-Left) region of the bacterial chromosome and the physiological significance of the characteristic early-growth-phase-dependent *fis* expression profile. This was achieved by exploiting the chromosome location, promoter and expression pattern of the *dps* gene. This gene is located in the Right macrodomain, where it encodes a ferritin-like, DNA-protecting protein, Dps, which is produced in high concentrations in stationary phase cells (49–54). Although Dps is a DNA binding protein, it does not influence transcription (51, 55). The *dps* gene is transcribed on entry to stationary phase using RpoS, the stress- and-stationary-phase sigma factor of RNA polymerase, although this can be overridden during oxidative stress when *dps* is transcribed using RpoD in an OxyR- and IHF-dependent mechanism (56, 57). Fis represses *dps* transcription by RNA polymerase containing the RpoD sigma factor, but not RpoS (58). Thus, Fis and Dps production is normally associated with opposite ends of the growth cycle: the early exponential and the stationary phases, respectively (46). The *fis* and *dps* genes are both transcribed in the opposite sense to the direction of DNA replication in the Left and Right replichores, respectively (Fig. 1A).

**Figure 1:**
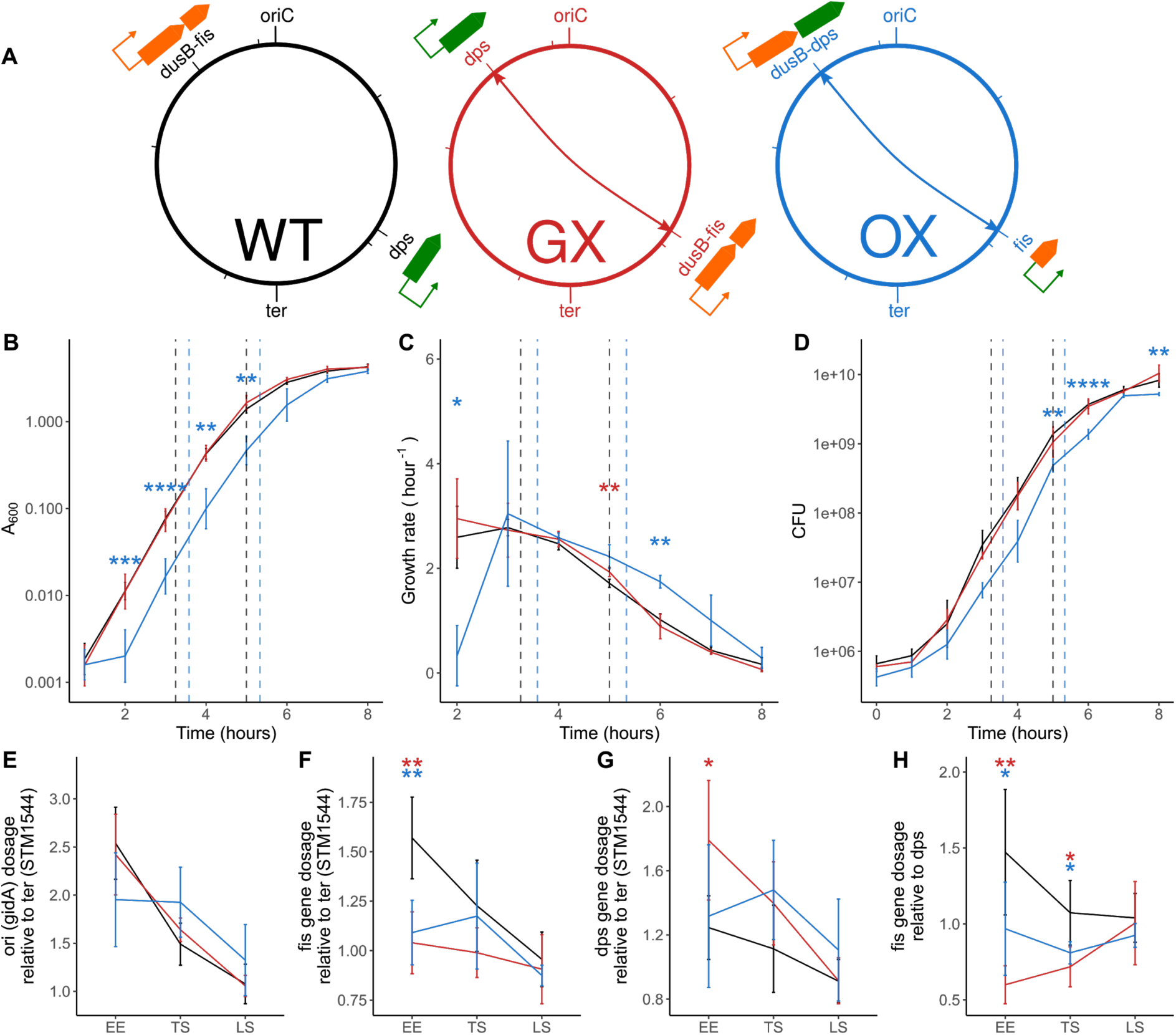
Genomic exchanges of *fis* and *dps* transcription units or ORFs. **(A)** Schematic depicting wildtype (WT) SL1344 strain with the original locations of *dusB-fis* and *dps* transcription units shown; the constructed Gene Exchange (GX) strain with the transcriptional units exchanged and the ORF Exchange (OX) strain with only the *fis-dps* ORFs exchanged. **(B)** Optical density based growth curves that show the impact of making these genetic changes on growth. Early Exponential (EE), Transition to Stationary (TS) and Late Stationary (LS) timepoints selected for other analyses are shown. Significance was determined relative to the wildtype using the Welch’s t-test and P-values were adjusted using BH method. Adjusted P-values < 0.05 are shown as *. **(C)** Growth rate curve derived from the OD based growth curves. **(D)** CFU based growth curves. **(E)** Origin to terminus gradients in these strains during the course of the growth curve calculated by assaying the abundance of *gidA* (*oriC* proximal gene) and *STM1544* (*ter* proximal gene) DNA using qPCR. Late stationary (LS) phase is the 24 hour time-point in the growth curve not shown in the earlier plots. **(F)** Dosages of the *fis* gene relative to the terminus during the course of the growth curve. **(G)** Dosages of the *dps* gene relative to the terminus. **(H)** Dosages of the *fis* gene relative to *dps* to understand the impact of gene position swap.

We anticipated that altering the expression pattern of the *fis* gene in a radical way, that falls short of eliminating Fis protein production completely, would provide valuable insights into 1. the robustness or the vulnerability of bacterial physiology in the face of changes to the expression of a prominent NAP gene 2. the impact on the operations of the cell of expressing the *fis* gene from a location in the opposite replichore to normal 3. the significance of the temporal expression pattern of the *fis* gene for the *in vivo* role of the Fis protein. To test our ideas, we relocated the *fis* gene, with its native control elements, from the NS-Left region of the chromosome of *Salmonella enterica* serovar Typhimurium (called *S*. Typhimurium from now on) to the *dps* locus in the Right macrodomain. In a second genetic exchange, just the protein-encoding, open reading frame of *fis* was substituted for that of *dps*, giving *fis* the stationary-phase-specific expression profile of the *dps* gene.

## RESULTS

### Reciprocally exchanging the *fis* and *dps* genes and open reading frames

The *fis* gene is the downstream component of the bicistronic *dusB-fis* operon (Fig. 1A). We repositioned the *Salmonella fis* gene’s transcription unit or its protein coding sequence to give them, respectively, just the geographical location (GX) or both the location and the growth-phase-dependent expression pattern of the *dps* gene (OX). In each case, the reciprocal exchange was made with the *dps* transcription unit (GX) or just with its protein coding region (OX). Since in GX the complete *dusB-fis* operon was moved to the *dps* site and the complete dps gene was relocated to the position on the chromosome normally occupied by *dusB-fis*, the native transcriptional and translational controls of *fis* and *dps* were preserved, with just the locations of the transcription units on the chromosome being altered (Fig. 1A). The construction of the GX and OX strains iis described in the Materials and Methods and Fig. S1. The genetic map locations of *fis* and *dps* in the wild type (WT), GX and OX strains are summarized in Fig. 1A. We used whole genome sequencing to rule out the presence of mutations that may have been introduced during strain construction (Fig. S2).

The complete gene exchange of the *dusB-fis* operon and *dps* did not produce measurable changes in the growth characteristics of the bacterium. The GX and WT strains exhibited similar growth profiles in liquid medium measured using absorbance (Fig. 1B), growth rate (Fig. 1C) or by counting colony-forming units (Fig. 1D). In contrast, the OX strain under-performed in comparison with the WT and GX in each of the three growth assessments with evidence of an extended lag phase being detected (Fig. 1 B, C, D). An exception to this was seen in the case of the growth rate of OX following the transition from exponential growth to stationary phase: here the growth rate of OX declined more slowly than that of the other two strains (Fig. 1C).

The locations of the *fis* and *dps* genes on the wild type chromosome are almost diametrically opposite, with *fis* being much closer to the origin of chromosome replication (*oriC*) and *dps* being nearer to the terminus, *ter* (Fig. 1A). The gene dosage gradients from *oriC* to *ter* were measured for each of the three strains using quantitative PCR (qPCR) of an origin-proximal gene (*gidA*) and a terminus-proximal one STM1544. The measurements were performed at three stages of growth: early exponential phase (EE), the transition from exponential growth to stationary phase (TS) and late stationary phase (LS) (for A_600_ values see materials and methods). The WT and GX strains had almost identical profiles while OX had a gradient with a higher variance at each stage of growth than the other two (Fig. 1E). The gene dosage of *fis* was lower in GX and OX in EE than in WT, as expected given its greater proximity to the terminus in the two exchanged strains (Fig. 1F). At EE, the *dps* gene dosage was greater in GX than in WT, as expected, but not greater in OX (Fig. 1G), possibly due to the lower growth rate and gene dosage gradient in the OX strain (Fig. 1C) stemming from the lower growth rate of the OX strain (Fig. 1C) at the early time point (Fig. 1E). At EE, the *fis* gene in the GX strain had the lowest gene dosage when measured relative to *dps* (Fig. 1H), thus clearly reflecting the *fis*-*dps* gene position exchange and associated *fis-dps* gene dosage exchange.

### Result of genetic exchanges on *fis* and *dps* gene expression and protein levels

The transcription of the *fis* (Fig. 2A) and *dps* (Fig. 2B) genes was measured by qPCR using the *hemX* gene as a benchmark. In the WT, each gene was expressed as expected with *fis* transcription being maximal in early exponential growth (Fig. 2A) and *dps* being transcribed maximally in stationary phase (Fig. 2B). In the GX strain, both *fis* and *dps* retained their overall expression profiles as seen in the WT but the levels of the *dps* transcript were higher (Fig. 2A, B). In the OX strain, the transcription profile of *fis* was the reverse of that seen in the WT: now *fis* transcript levels were highest in stationary phase and lowest in early exponential phase (Fig. 2A). In contrast, *dps* expression was highest in exponential growth and lowest in stationary phase, the reverse of the WT pattern (Fig. 2B). Western blot analyses showed that Fis (Fig. 2C) and Dps (Fig. 2D) production patterns resembled the transcription patterns for the corresponding genes: Dps production in OX mimicked Fis production in the WT while Fis was produced in OX in a pattern that resembled that of Dps in the WT. Notably, the expression of *fis* and *dps* genes determined using RNA-seq (Fig. 2E, F) matched the expression determined using qPCR (Fig. 2A, B). These data showed that relocating the *dusB-fis* operon or the *dps* gene had a negligible effect on their expression patterns or on the production profiles of their products. However, exchanging just the *fis* and *dps* protein coding regions, with the concomitant switching of cognate regulatory elements, produced reciprocal changes in gene expression and protein production. We next examined the impacts of these changes on Fis protein binding patterns throughout the genome, on the downstream expression of the Fis regulon and on the physiology of the bacterium.

**Figure 2:**
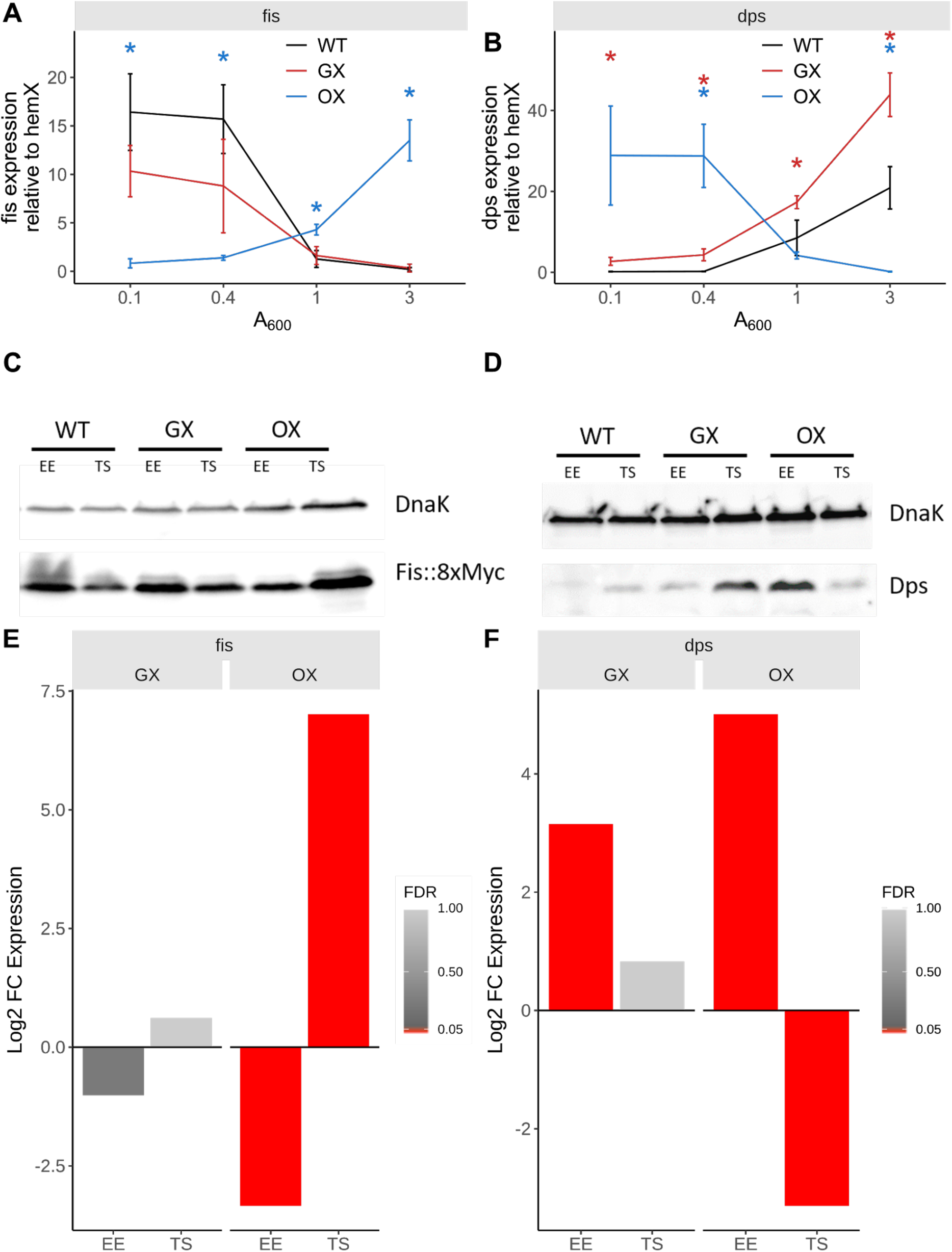
Gene expression changes. **(A)** Expression of *fis* transcript during the course of the growth curve assayed using RT qPCR. Significance was determined relative to the wildtype using the Welch’s t-test and P-values were adjusted using the BH method. Adjusted P-values < 0.05 are shown as *. WT (black), GX (red), OX (blue). **(B)** Expression of *dps* transcript during the growth curve. WT (black), GX (red), OX (blue). **(C)** Fis protein levels determined by immunoblotting with anti-Fis monoclonal antibody. **(D)** Dps protein levels determined by immunoblotting with anti-Dps polyclonal serum. **(E-F)** Expression changes in *fis* and *dps* genes determined by RNA-seq in the GX and OX strains at EE and TS phases of growth. Colour bar indicates FDR and bars representing genes with FDR < 0.05 are red.

### Fis binding to the genome in the GX and OX strains

ChIP-seq was used to examine the intensity and the distribution of Fis protein binding to the genomes in the GX and OX strains compared to the WT (Fig. 3 & S3, Fis motif: Fig. S4). In both the GX and OX strains, the median Fis binding intensity was lower than in the WT (Fig. 3A). The intensity of Fis binding changed over a greater range, and was significantly changed at more Fis peaks, in the OX than in the GX strain (Fig. 3A, 3C). Changes in Fis binding intensity in the GX strain positively correlated with those in the OX strain suggesting these two strains experienced similar changes in Fis binding across genomic loci (Fig. 3B, 3C). Binding intensity changes showed a geographical distribution in the OX:GX comparison (Fig. 3C). On average in both OX and GX, the reduction in Fis binding was greater around the chromosome origin than around the terminus. However, the magnitude of the change was greater for OX:WT than for GX:WT. In both OX and GX, the Fis protein is being produced from a locus that is closer to the terminus than to the origin (i.e. the gene location is diametrically opposed to that in the WT) (Fig. 1A). We also examined the binding patterns of Fis at the negatively autoregulated *dusB-fis* promoter in all three strains. OX and the WT had very similar patterns of Fis binding peaks but the peaks showed a reduced amplitude in GX (Fig. S6). This may reflect the influence of the new chromosomal neighborhood following the relocation of those Fis binding sites together with *dusB-fis* to a novel chromosomal site (the *dps* locus) in GX.

**Figure 3:**
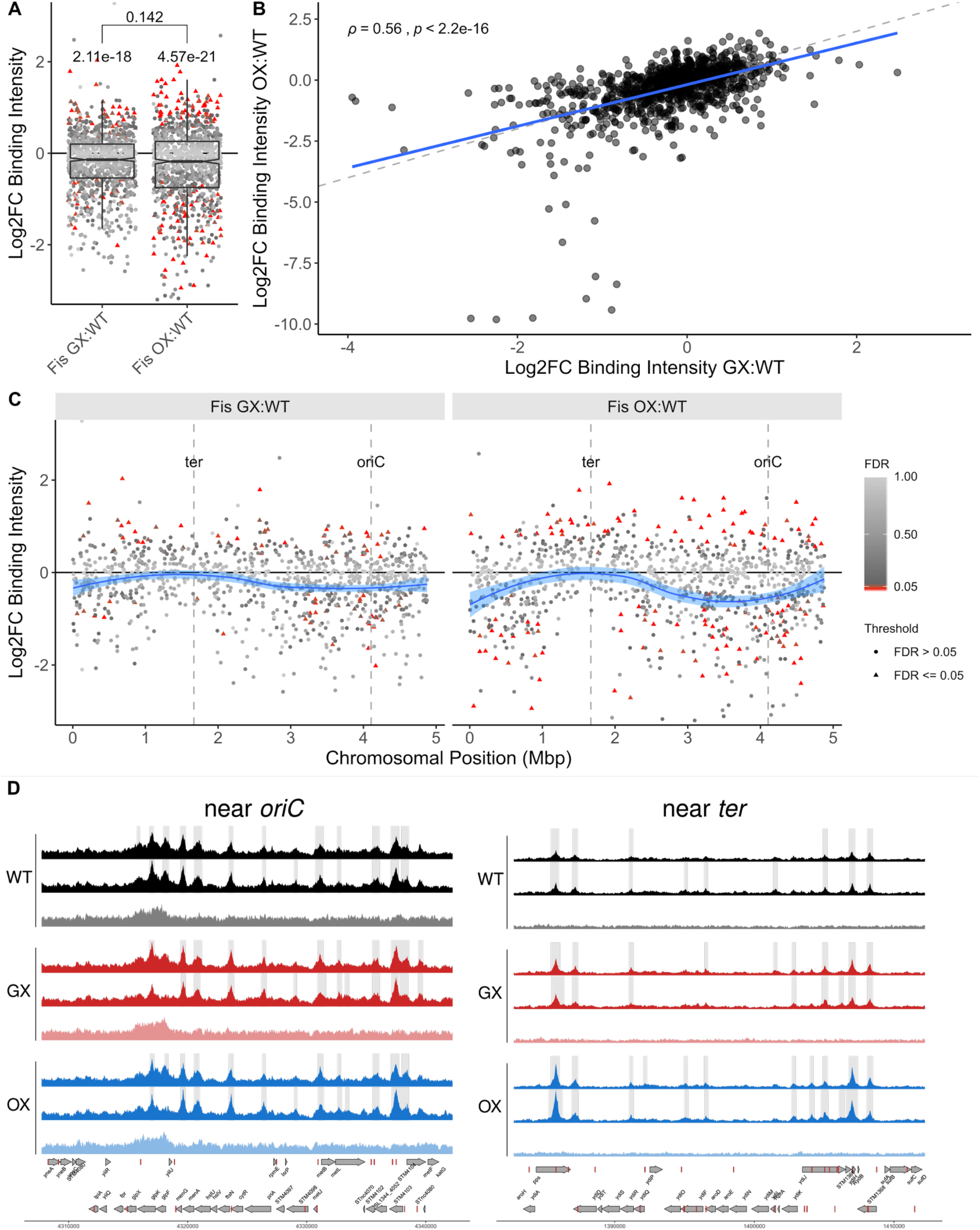
Fis binding changes assayed by ChIP-seq during EE phase. **(A)** ChIP-seq was used to identify Fis peaks in WT, GX and OX. The R package DiffBind was then used to find Log_2_ fold change in binding intensities of Fis in these peaks in the GX and OX strains relative to the wild type. Significant peak intensity changes (FDR < 0.05) are shown as red filled triangles. To determine if the medians of distributions of Log_2_ fold change in binding intensities differed significantly from 0 (no change in Fis binding relative to WT), the Wilcoxon rank sum test was performed with µ = 0 and the P-values are displayed over the individual distributions. The difference between the medians of the two, GX:WT and OX:WT, distributions was not significant with the Wilcoxon rank sum test and is shown on the bracket above the single sample tests (using the less conservative Welch’s t-test here gives P value = 0.0105). **(B)** Spearman’s correlation of peak intensity changes in the GX with the OX strain. Dashed line has slope of 1 and indicates the position where the fold changes in Fis binding intensity in GX would be equal to those in OX. Blue solid line is the regression linear fit to the data. **(C)** Peak intensity changes along the chromosomal position shown in millions of base pairs. The blue curve shows the local regression peak intensity change calculated using the R loess function. In calculating these intensity changes, the appropriate mocks (WT mock for WT peaks, GX mock for GX peaks, OX mock for OX peaks) were used to control for *ori-ter* gradient changes (Fig. S5). **(D)** Fis ChIP-seq coverages of the WT, GX and OX strains (two biological replicates and mock) of example regions near *oriC* and *ter* loci. Grey shaded regions show peaks called in each sample relative to the mock. Genes (grey arrows) and Fis binding motifs (red blocks) are shown on the + and - reference strands. ChIP-seq data for plasmids can be found in Fig. S3.

*Salmonella* Typhimurium SL1344 harbors two large plasmids: pSLT (the virulence plasmid) and pCol1B9, both of which are bound by Fis (Fig. S7). As with the chromosome, both plasmids displayed reduced Fis binding at most loci in the GX and OX strains (Fig. S7A) and these changes were positively correlated between the two strains (Fig. S7B).

### The impact of the genetic exchanges on Fis target gene expression

RNA-seq was used to compare the transcriptomes of the GX and the OX strains with the WT in cultures at the early exponential phase and at the exponential-to-stationary transition phase of the growth cycle (Fig. 4). OX showed a higher number of genes with altered expression than GX at both phases of growth (Fig. 4A) and the changes in expression were also greater in OX than in GX (Fig. 4B). Furthermore, the difference between the median log_2_ fold change in expression (log_2_FC) of the Fis target genes and the non-target genes was more significant in the OX strain compared to the GX. In the OX strain, the median log_2_FC of the Fis target genes was lower in the early exponential phase, and greater in the transition to stationary phase, than that of the non-target genes. This difference in Fis target and non-target gene expression mirrors the greater Fis levels during exponential phase and lower Fis levels in the transition to stationary phase in the OX strain. As with the Fis binding data, we saw slight positive correlations between the GX and OX strains at both EE and TS phases (Fig. 4C), thus highlighting similarity in the Fis binding changes and associated gene expression changes in the two strains. When the data were plotted as a function of position along the chromosome, the OX strain was again found to have a far greater degree of disturbance to normal expression of its transcriptome, compared to the WT, than GX and differentially expressed genes were scattered throughout the chromosome (Fig. 4D). These global gene expression patterns were consistent with the much greater changes seen to *fis* expression, Fis protein production (Fig. 2) and Fis protein binding in the OX strain (Fig. 3).

**Figure 4:**
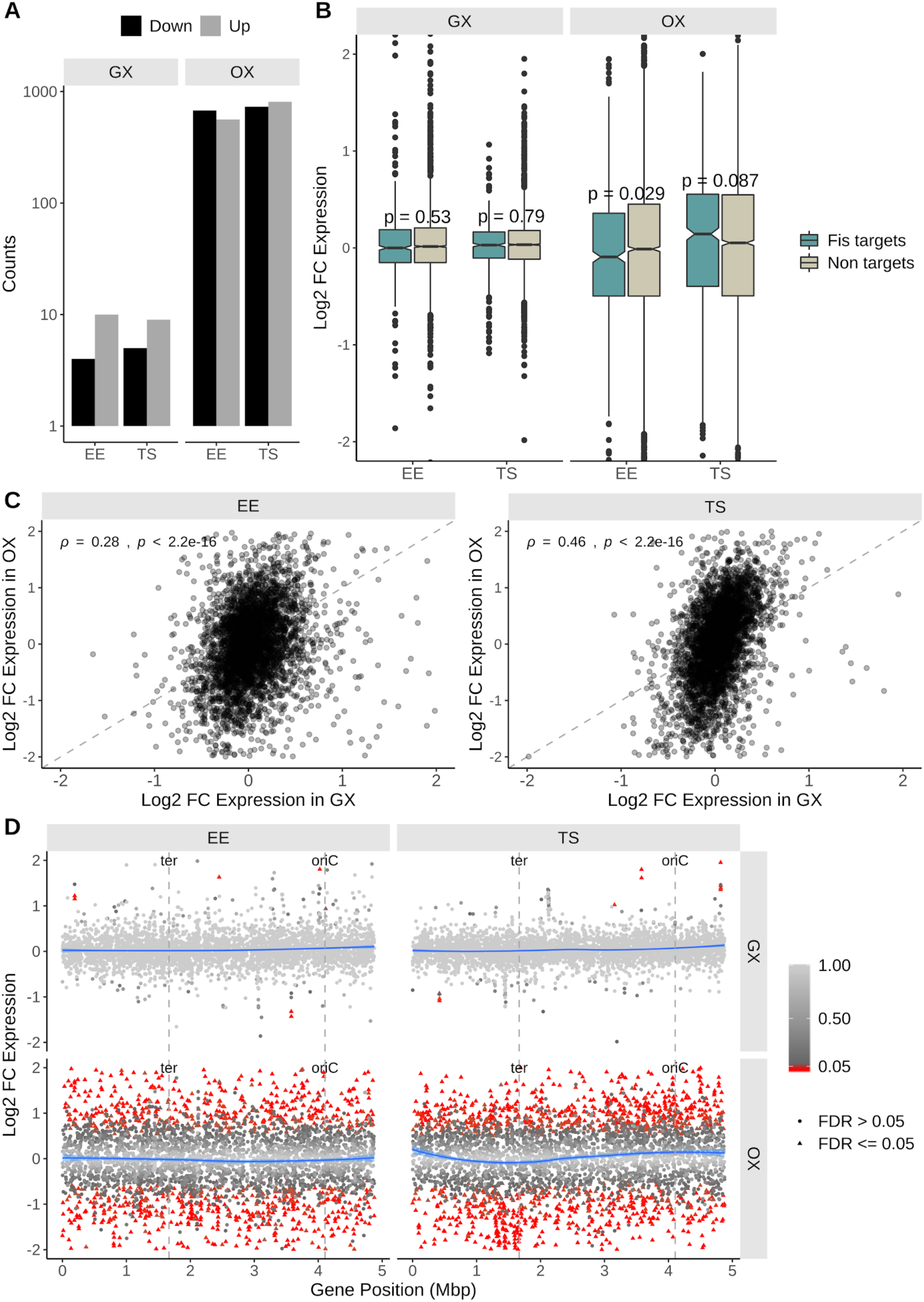
Transcriptomic changes assayed by RNA-seq. **(A)** Counts of differentially expressed genes. **(B)** Box plot distribution representations of Log_2_ fold changes in expression of genes in the GX and OX strains. Genes have been split into two groups based on whether or not they are predicted to be Fis targets. P values obtained using the two sample Wilcoxon test for the comparison between Fis targets and non targets is mentioned over each pair of box plots. **(C)** Spearman’s correlation of Log_2_ fold changes in expression of genes in the GX with the OX strain. Dashed line has slope of 1 and indicates the position where the fold changes are equal. **(D)** Log_2_ fold changes in expression of genes along their position on the chromosome in millions of base pairs. The blue curve shows the local regression peak intensity change calculated using the R loess function. Significant gene expression changes (FDR < 0.05) are shown as red filled triangles.

Among known Fis-dependent genes, those encoding the three type 3 secretion systems found in *Salmonella* exhibited changes in expression (Fig. S8). These genes encode the flagellar apparatus and the SPI-1 and SPI-2 injectisomes. In each case, the largest, statistically significant effects were seen in the OX strain (Fig. S8). The altered patterns of transcription detected in SPI-1 and SPI-2 in the RNA-seq experiments were investigated in more detail using *gfp* reporter gene fusions to representative promoters from each pathogenicity island: P_*prgH*_ (SPI-1) and P_*ssaG*_ (SPI-2) (Fig. 5A, 5B). These experiments confirmed that both promoters were dependent on Fis because their full activities were lost in a *fis* knockout mutant. In contrast, the complete removal of Dps from the cell by introducing a *dps* knockout mutation had no effect on either promoter (Fig. 5B). In OX, the SPI-1 promoter P_*prgH*_ had reduced activity compared to WT at 5 h in batch culture but was significantly more active at 10 h (Fig. 5A, 5B). P_*ssaG*_ from SPI-2 was also less active in OX at earlier stages of growth but showed no significant difference to the WT at 20 h (Fig. 5A, 5B). The P_*prgH*_ promoter was as active in the GX strain as it was in the WT at all stages of growth; the P_*ssaG*_ promoter differed from the WT by being less active at the 7.5-h time point.

**Figure 5:**
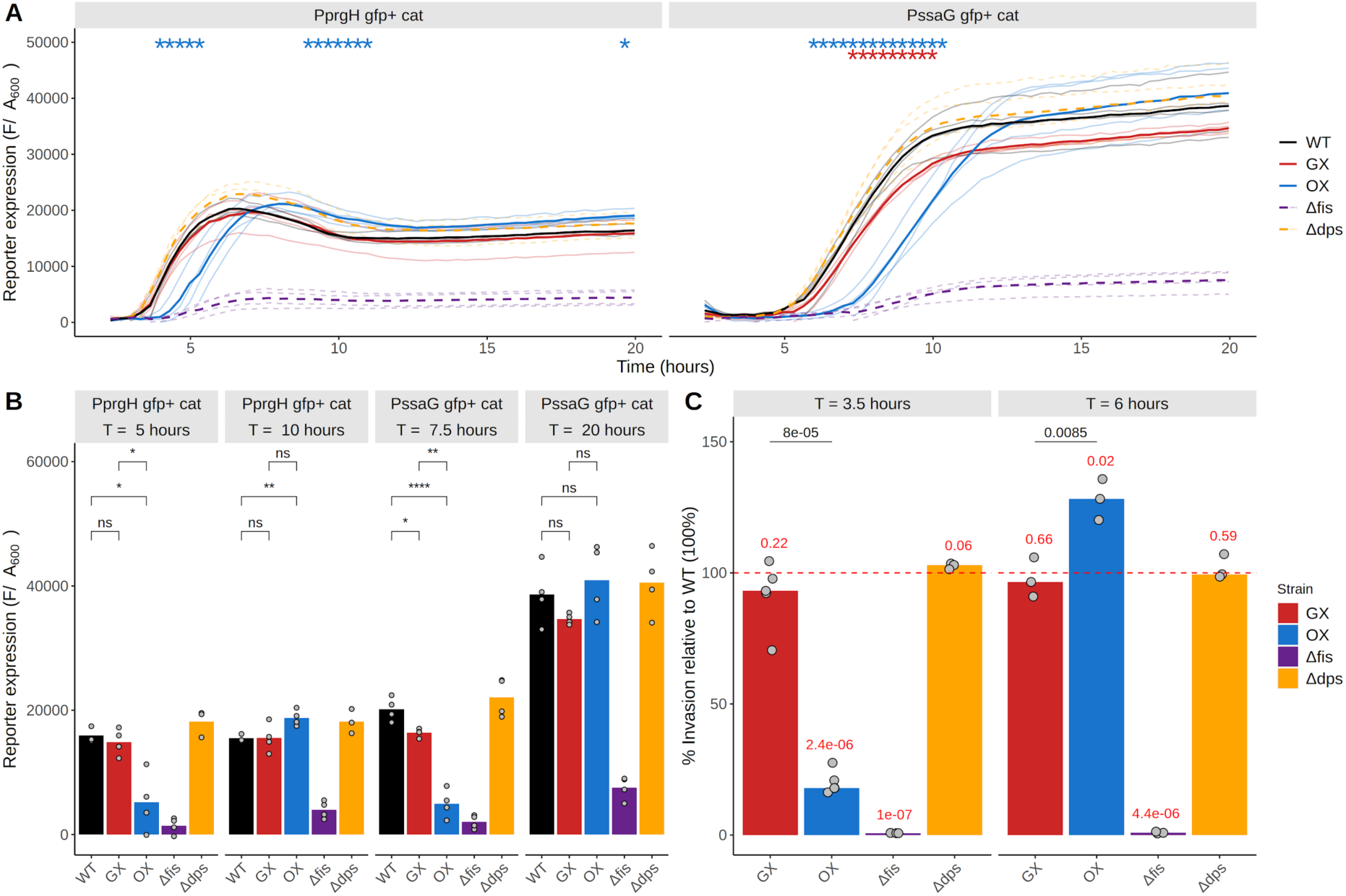
Pathogenicity island expression changes assayed using chromosomal *Pgfp*^*+*^ reporters. **(A)** SPI-1 expression was reported by the P_*prgH*_ *gfp*^*+*^ and SPI-2 expression was reported by the P_*ssaG*_ *gfp*^*+*^ chromosomal fusions. Reporter expression is measured by fluorescence (F) of *gfp*^+^ expression divided by A_600_. Mean of biological replicates are shown as solid lines, and the individual biological replicates are shown in the background as the lighter shaded lines. Significance was determined relative to the wildtype using the Welch’s t-test and P-values were adjusted using the BH method. Adjusted P-values < 0.05 (WT vs GX and WT vs OX comparisons only) are shown as *. **(B)** A slice of the data in (A) showing reporter expression at early (SPI-1: 5 hours; SPI-2: 15 hours) vs. later (SPI-1: 15 hours; SPI-2: 20 hours). Bars represent median value and the individual biological replicates are shown by the scatter. **(C)** Results of epithelial cell invasion assay carried out with subcultures grown to two different timepoints (early: 3.5 hours; late: 6 hours). Plotted are percentage invasions of the strains relative to the WT (100%, not shown, depicted by the red dashed line) in HeLa cells determined 90 minutes post infection. Median values of % invasion are represented by the bars and values of biological replicates are shown by the scatter. P values obtained from one sample t-tests (µ = 100) are shown in red above each strain. P values obtained from the Welch’s two sample t-tests comparing between GX and OX are mentioned over these comparisons in black.

Given the changes to SPI-1 and SPI-2 gene expression detected in the OX strain, this strain and the GX and WT strains were compared in an *in vitro* infection assay using cultured HeLa cells. The OX strain exhibited a strong reduction in infectivity at the 3.5-h time point, although not as strong as the *fis* knockout mutant control (Fig. 5C). In contrast, the GX strain showed a modest reduction in invasiveness compared to the WT at both the 3.5- and the 6-h time points. The OX strain had enhanced invasiveness at 6 h (Fig. 5C). The *dps* knockout mutant was indistinguishable from the wild type at either time point.

We also examined the transcriptomes of the two large plasmids in GX, OX and the WT (Fig. S9). As with the chromosome, genes on the pSLT and pCol1B9 plasmids had shown changes in expression in the OX strain, but not GX strain, compared with the WT, showing that the changes made to Fis production in OX had impacts beyond the chromosome (Fig. S9). Notably, in the OX strain at TS phase, we were able to see an upregulation of the *tra* genes on pCol1B9.

## DISCUSSION

We have investigated the effect of repositioning the *fis* gene on the *S.* Typhimurium chromosome. This gene encodes the conditionally abundant nucleoid-associated protein, Fis, which is present in over 50,000 copies in rapidly growing bacteria but with very few copies in stationary phase cells (35). Previous work investigating the effects of regulatory gene translocation has used *lacI*, the gene encoding the transcriptional repressor, LacI, present in approximately 40 monomers per cell (6, 59). Fis has a much larger regulon than LacI, influencing the transcription of hundreds of genes, positively or negatively (19), and it plays a role in organizing the local structure of the nucleoid (60, 61). Its influence extends beyond transcription and includes effects on chromosome replication (26), transposition (28) and site-specific recombination (20, 22). We sought to investigate if, due to the pervasive influence of Fis, altering the location of the *fis* gene might produce significant changes to bacterial physiology. We were encouraged in our investigation by a previous study in which the translocation of the *dusB-fis* operon to the Ter region of the *E. coli* chromosome correlated with changes to the topology of a reporter plasmid and stress resistance, albeit without alterations to Fis protein levels in the cell (12). This lack of alteration to Fis protein levels was a consequence of an increase in *fis* expression, which compensated for the reduction in *fis* gene copy number (12). In our study, the same effect was not observed in the GX strain: *fis* expression and Fis concentration remained at similar levels to those of the WT strain in EE, despite the reduced *fis* gene dosage at this point in growth. Furthermore, the GX strain did not exhibit a growth defect whereas the *E. coli* strain with a translocated *dusB-fis* operon described by Gerganova *et al*. did (12). Together, these data indicate that other factors (including bacterial species differences) affect *fis* expression at a given location on the bacterial chromosome. In addition to *dusB-fis* translocation, in our experiments we also studied the effects of rewiring the *fis* gene so that it had the expression profile of *dps*, a gene with a growth phase expression profile that is diametrically opposed to that of *fis* (46, 58). By doing so we were able to saturate the cell with Fis in the stationary phase of growth, a period when Fis is normally undetectable. Similarly, we could reduce the production of Fis to a very low level in EE (early exponential) phase, an interval when the protein is normally very abundant.

DNA-binding regulators of transcription and of other molecular processes must communicate with their genomic targets by translocation through the cell and some studies based on the LacI protein suggest that the distance to be travelled influences the efficiency of the regulatory connection (62–64). There is some evidence that translated mRNA is spatially constrained in bacteria, remaining close to the site of transcription (65) while other work suggests that the localisation of mRNA is driven by the nature of the protein product and its cellular location (66, 67). Collectively, these studies suggest that the physical location of a gene in the folded chromosome may determine the influence exerted by its protein product in the cell. A corollary to this proposition is that the repositioning of a gene may alter the sphere of influence of its product.

Our results with the GX strain show that moving the entire *dusB-fis* operon to the Ter-proximal site normally occupied by the *dps* gene has relatively modest impacts on *Salmonella* physiology. This was true in terms of growth kinetics of GX compared with the wild type (Fig. 1B-D), in the transcriptomic comparison (Fig. 4) and when GX and the wild type were compared in terms of SPI-1 *prgH* virulence gene expression (Fig. 5A, 5B). In contrast, the activity of the SPI-2 virulence gene promoter P_*ssaG*_ was significantly reduced in GX compared to the wild type and GX exhibited a reduction in HeLa cell invasion (Fig. 5). Both the P_*prgH*_ and the P_*ssaG*_ promoters were shown previously to have reduced activity in a *fis* knockout mutant of *Salmonella*, indicating a dependence on this NAP for normal function (19). Our new data show that altering the physical location of the *fis* gene influences P_*ssaG*_ negatively and is associated with a reduction in a virulence phenotype. In GX, the *dusB-fis* operon is moved from the Non-structured Left region of the chromosome, where SPI-1 is also located, to the Right macrodomain, adjacent to the right-hand segment of the Ter macrodomain where SPI-2 is found. The production of the Fis protein in GX had increased at the transition to the stationary phase of growth compared to the wild type (Fig. 2C), perhaps reflecting the homeostatic resetting of Fis levels reported previously when the *dusB-fis* operon was moved to a Ter-proximal site in *E. coli* (12). Looking in detail at the SPI-1 and SPI-2 pathogenicity islands, the SPI-2 gene cluster shows the greatest decrease in expression in the OX strain at the onset of stationary phase (Fig. S8) which is consistent with the impact on virulence (Fig. 5C).

Overall, moving the *fis* gene to the native chromosomal location of *dps* and expressing *fis* using the regulatory elements normally used by *dps* in the OX strain had much more profound effects on *Salmonella* physiology than those seen in the GX strain. These effects included an extended lag phase in the OX strain when growing in batch culture (Fig. 1), a much more profound impact on the transcriptome with many more genes in OX being affected than in GX and with a greater amplitude of positive and negative changes in transcription (Fig. 4) and a greater influence of the expression of key virulence genes and of a virulent phenotype (Fig. 5). The OX strain produced the Fis protein with a profile equivalent to that of Dps in the wild type and expressed Dps in a Fis-like manner (Fig. 2). Dps has a negligible impact on transcription (51) so the impact on gene expression in OX is likely to reflect differences in Fis protein levels, a proposition that is supported by the finding that so many known Fis targets are among the genes showing altered expression.

Can the changes to *Salmonella* physiology in the GX and OX strains be explained by novel patterns of Fis distribution along the chromosome? The ensemble experiments conducted in this study relied on ChIP-seq to monitor Fis protein distributions during the early exponential phase of growth in living cells. The intensities of the protein binding peaks did change in the two rewired strains, especially in the vicinity of Ter, with OX showing by far the greater change compared to the wild type (Fig. 3C, 3D). This likely reflects the more radically altered pattern of Fis production in OX, where the transcription signals of the *dps* gene are driving the expression of *fis* (Fig. 1A). The bacterium used in this study, *S*. Typhimurium strain SL1344, also harbours the large plasmids pSLT (a virulence plasmid) and pCol1B9. These autonomously replicating circular DNA molecules are independent of the chromosome and they also contain Fis binding sites. The patterns of Fis binding to the two plasmids were generally similar in the GX, OX and wild type strains (Fig. S3). Plasmid gene expression was unaffected in the GX strain compared to the wild type but statistically significant differences were detected in OX (Fig. S9).

With the exception of H-NS (68, 69), no NAP in *Salmonella* is essential for the life of the bacterium and this includes Fis and Dps. The contributions made by NAPs seem to be auxiliary to the functions that they influence. Our findings show that the *Salmonella* genome is well buffered against even quite radical changes to the production of Fis and Dps. The reciprocal inversion of their growth-phase-dependent production profiles, so that Fis assumes the pattern that is characteristic of Dps, and *vice versa*, does not seriously perturb the life of the cell, although the OX strain does exhibit the many physiological changes that have been documented here. Prominent among these are the impacts we detected on *Salmonella* plasmid and virulence gene expression and pathogenesis. The inversion of the SPI expression pattern in the OX strain relative to the WT (Fig. S8B), as well as the inversion of the pattern of invasion of this strain relative to the WT (Fig. 5C), demonstrates the role of the timing of Fis expression in controlling the normal timing of expression of these islands. These changes are likely to have their most telling effects when the bacterium is experiencing stress or when trying to compete with the wild type, compromising its ability to compete with the wild type strain.

## MATERIALS AND METHODS

### Bacterial strains, growth media and chemicals

Bacterial strains used in this study, their genotypes and source, where applicable, are listed in Table S1; all strains are derivatives of *Salmonella enterica* serovar Typhimurium strain SL1344. Unless otherwise stated bacterial strains were cultured in Miller Lysogeny broth (LB, 1% NaCl). Kanamycin, carbenicillin/ampicillin and chloramphenicol were used at concentrations of 50 µg/ml, 100 µg/ml and 35 µg/ml, respectively.

### Strain construction

All strains generated during this study were constructed using phage λ Red recombinase mediated homologous recombination (70) and P22 transduction (summary schematic in Fig. S1). To construct the <u>G</u>ene e<u>x</u>change (GX) and <u>O</u>pen reading frame e<u>x</u>change (OX) strains, first, the FRT flanked kanamycin resistance gene (*kan*), amplified from pKD4, was inserted 62 base pairs (bp) downstream of the *fis* ORF (SL1344 *fis::kan*) and the FRT flanked chloramphenicol resistance gene (*cat*), amplified from pKD3, was inserted 190 bp upstream of the *dps* ORF (SL1344 *dps::cat*). These strains were used as templates to amplify either *fis::kan* ORF or the *dusB-fis::kan* and *dps::cat* transcriptional units. SLiCE (71) was used to generate a *dusB-dps-cat* construct on the plasmid pBluescript SK(+). This plasmid was used as a template to amplify the *dusB-dps-cat* transcriptional unit (where the *fis* ORF is replaced by the *dps* ORF). As previously described, these amplicons were used to replace the reciprocal gene, generating the 2x intermediate strains in the process that harboured two copies of either the *fis* or *dps* transcriptional units or ORFs (Table S1, Fig. S1). P22 phage mediated generalized transduction was used to generate the GX and OX strains. Finally, the FRT flanked antibiotic resistance genes were eliminated using a helper plasmid (pCP20) encoding FLP recombinase.

Strain variants of SL1344 and the exchange strains expressing 8X Myc-epitope-tagged Fis were constructed using a modified version of phage λ Red recombinase mediated recombination (70).

The DNA oligonucleotides used in this study are listed in Table S2.

### Bacterial growth measurement

Bacterial cell density was measured using absorbance and viable counts.

Absorbance was measured, using the Thermo Scientific BioMate 3S spectrophotometer, at 600 nm (A_600_) using sterile LB for blank correction. Cell densities with A_600_ greater than 1 were diluted 1:10 in sterile LB before measurement.

Viable counts were performed by plating serial ten fold dilutions of the culture made in sterile phosphate buffered saline (PBS) on sterile LB agar plates. Colonies obtained after overnight incubation at 37 °C were counted and multiplied by the appropriate dilution factors to give the viable cell density in terms of colony forming units (CFU).

Overnight cultures were grown for at least 16 h in 4 ml LB in tubes at an angle at 37 °C and 200 rpm orbital shaking. For performing growth experiments, overnight cultures were diluted to A_600_ of 0.003 in 25 ml LB in 250 ml conical flasks. These were incubated at 37 °C with 200 rpm orbital shaking.

For all experiments bacterial cultures were harvested at the early exponential (EE, A_600_ of 0.4), transition to stationary (TS, A_600_ of 3) or late stationary (LS, A_600_ of 3.5, 24 h from inoculation) phases of growth.

### Estimating gene dosages using quantitative reverse transcription PCR (RT-qPCR)

Bacterial chromosomal DNA was isolated by harvesting 1 ml of cultures grown to EE, TS and LS phases of growth by centrifugation at 16000 × g for 1 min. The cell pellet was resuspended in sterile distilled water and boiled at 100 °C for 5 min, vortexed and cell debris was pelleted by centrifugation again. The supernatant was mixed with 1 volume of chloroform, and mixed by vortexing. The upper aqueous layer was isolated by centrifugation at 16000 × g for 10 min. The concentration was determined using a DeNovix DS-11 spectrophotometer using A_260_ and adjusted to 100 ng/µl with nuclease free water.

The gDNA was probed using primers specific to target genes in the StepOnePlus Real Time PCR system. Each 20 µl reaction, set up in the wells of a 96 well plate, contained 1x FastStart Universal SYBR Green Master (Rox) (Merck, Wicklow, Ireland), 0.6 µM primer pair, 8 µl gDNA and sterile distilled water to 20 µl. For each primer pair a standard curve of ten-fold serially diluted SL1344 gDNA was included. Threshold cycle (Ct) values of gDNA samples were checked against the standard curves to ensure that they fell in the linear range for each primer pair and the concentration of gDNA in the samples were estimated from the standard curve. Cycle conditions for qPCR reactions were as follows; 95 °C for 10 min, followed by 40 cycles of 95 °C for 15 s, 60 °C 1 min.

Oligonucleotide primers were designed against an *oriC* (*gidA*) proximal gene and a Ter proximal gene (*STM1554*). To determine the gene dosage gradient, the ratio of *oriC*-proximal DNA to Ter-proximal DNA was identified. Specific gene dosages of *fis* and *dps* were determined by comparing the quantities of these genes to *STM1554.*

### Estimating gene expression changes using quantitative reverse transcription PCR (RT-qPCR)

At optical densities of 0.1, 0.4, 1 and 3, further growth was halted and intracellular RNA was stabilised by the addition of 0.4 volume stop solution (5% v/v phenol, pH 4.3, in ethanol) and incubation on ice for 30 min. A volume of cells equivalent to a cell density of 1 ml culture with A_600_ of 1, were harvested. Cells were pelleted by centrifugation at 3220 × g for 10 min at 4 °C. Cells were resuspended in Tris EDTA buffer (TE, pH 8.0) containing 50 mg/ml lysozyme. RNA was extracted using the SV Total RNA Isolation kit (Promega, Wisconsin, USA). RNA was DNase treated using the Turbo DNase kit (Invitrogen). RNA integrity was assessed on a HT gel (72) and quantified using the DeNovix DS-11 spectrophotometer using A_260_.

RNA (400 ng) was reverse transcribed into cDNA using random oligos and the GoScript Reverse Transcriptase kit (Promega). The cDNA was probed using primers specific to target genes in the StepOnePlus Real Time PCR system as described above. The *hemX* gene was used as a control gene (assumed unchanged expression) and the expression changes of the other genes were calculated against *hemX*.

Oligonucleotide primer pairs used in qPCR are listed in Table S3.

### Estimating protein levels using immunoblotting

At the EE or TS phases of growth, a volume of cells equivalent to a cell density of 1 ml culture with A_600_ of 1, were harvested and resuspended in 350 µl PBS and transferred to a sonication tube. Cells were lysed by sonication with 10 cycles, with each cycle consisting of 10 seconds bursts (10 µm amplitude) followed by 30 seconds incubation on ice. Samples were diluted with equal volume 2x Laemmli sample buffer (Final concentration: 4% SDS, 20% glycerol, 10% 2-mercaptoethanol, 0.004% bromophenol blue, 0.125 M Tris HCl, pH 6.8) and heated at 100 °C for 5 min prior to loading on a 12.5% SDS PAGE gel.

Proteins were separated by electrophoresis in running buffer (25 mM Tris-HCl, 190 mM Glycine, 0.1% (w/v) SDS), at 60 V through the stacking gel, which increased to 130 V as the dye moved through the resolving gel until the dye reached the base of the gel and then transferred onto methanol activated PVDF membranes (0.2 µm) in transfer buffer (20% (v/v) methanol, 25 mM Tris, 190 mM Glycine), at 300 mA for 1 h 45 min using a Trans-Blot apparatus (Bio-Rad).

Blocking of the membrane was performed for 1 h at room temperature with a blocking solution (5% skimmed milk powder, phosphate buffered saline, 0.1% Tween 20, PBST), while rocking. Membranes were probed overnight at 4°C using the following primary concentrations; 1:50000 Mouse-α-DnaK monoclonal antibody (Abcam, ab69617), 1:50000 Mouse-α-C-Myc monoclonal antibody (Sigma, M4439), 1:2000 Rabbit-α-Dps polyclonal serum (Prof. Regine Hengge, Humboldt Universitat zu Berlin). All antibodies were diluted to the appropriate concentration in PBST containing 5% BSA. Membranes were washed with PBST for 5 min each while rocking. HRP-conjugated secondary antibodies were diluted 1:10000 and 1:2000 for polyclonal Goat-α-Mouse (Bio-Rad, 170-6516) and Goat-α-Rabbit HRP (Dako, P0448), respectively. Secondary antibodies were diluted to the appropriate concentration in PBST containing 5% skimmed milk powder (Lab M). Membranes were incubated with secondary antibodies for 1 h 30 min at room temperature while rocking. Membranes were then washed three times with PBST and once with PBS. Blots were incubated in ECL reagent (Pierce) for 1 min, and bands were visualized using an ImageQuant LAS 4000 scanner (GE Healthcare).

### Whole Genome Sequencing (WGS) and Single Nucleotide Polymorphism (SNP) analysis to check constructed strains

#### DNA extraction

Chromosomal DNA for whole genome sequencing was prepared by a phenol-chloroform method which caused minimal shearing to DNA. Briefly, a 2 ml of an overnight culture was harvested by centrifugation at 16000 × g and the cell pellet was resuspended in TE pH 8.0 (100 mM Tris-HCl pH 8.0, 10 mM EDTA pH 8.0). Lysis was performed by the addition of 1% (w/v) SDS, 2 mg/ml proteinase K and incubating at 37 °C for 1 h. DNA was isolated by the addition of 1 volume of phenol pH 8.0: chloroform : isoamyl alcohol (25 : 24 : 1), followed by centrifugation at 16000 × g to separate the aqueous and organic layers. Following this, 1 volume of chloroform was added to the aqueous layer, and centrifugation was repeated. To remove contaminants 0.3 M sodium acetate pH 5.2 and 5 volumes of 100% (v/v) ethanol were added and precipitated at -20°C for 1 h. The DNA was pelleted by centrifugation, washed once with 70% (v/v) ethanol, dried at 65 °C for up to 5 min and resuspended in 50 µl sterile distilled water. Total gDNA was treated with 100 mg/ml RNase A at 37 °C for 30 min. Following this, DNA was isolated by the addition of 1 volume of phenol pH 8.0: chloroform : isoamyl alcohol as described above.

#### Library preparation

Library preparation and sequencing for all strains except OX and 2xdO was performed by MicrobesNG (http://www.microbesng.uk) which is supported by the BBSRC (grant number BB/L024209/1). DNA was quantified in triplicates with the Quantit dsDNA HS assay in an Eppendorf AF2200 plate reader. Genomic DNA libraries were prepared using Nextera XT Library Prep Kit (Illumina, San Diego, USA) following the manufacturer’s protocol with the following modifications: two nanograms of DNA instead of one were used as input, and PCR elongation time was increased to 1 min from 30 seconds. DNA quantification and library preparation were carried out on a Hamilton Microlab STAR automated liquid handling system. Pooled libraries were quantified using the Kapa Biosystems Library Quantification Kit for Illumina on a Roche light cycler 96 qPCR machine. Libraries were sequenced on the Illumina HiSeq using a 250bp paired end protocol. Reads were adapter trimmed using Trimmomatic 0.30 with a sliding window quality cutoff of Q15 (73). De novo assembly was performed on samples using SPAdes version 3.7 (74), and contigs were annotated using Prokka 1.11 (75).

For the OX and 2xdO strains, library preparation was performed by the Sanger Institute (Hinxton, UK.). Samples quantified with Biotium Accuclear Ultra high sensitivity dsDNA Quantitative kit using Mosquito LV liquid platform, Bravo WS and BMG FLUOstar Omega plate reader and cherrypicked to 200ng / 120ul using Tecan liquid handling platform. Cherrypicked plates were sheared to 450bp using a Covaris LE220 instrument. Post sheared samples were purified using Agencourt AMPure XP SPRI beads on Agilent Bravo WS. Library construction (ER, A-tailing and ligation) was performed using NEB Ultra II custom kit on an Agilent Bravo WS automation system. PCR was set-up using KapaHiFi Hot start mix and IDT 96 iPCR tag barcodes on Agilent Bravo WS automation system (PCR cycles, 6 standard cycles, 1) 95 °C 5 mins, 2) 98 °C 30 secs, 3) 65 °C 30 secs, 4) 72 °C 1 min, 5) Cycle from 2, 5 more times, 6) Incubate 72°C 10 mins. Post PCR plate purified using Agencourt AMPure XP SPRI beads on Beckman BioMek NX96 liquid handling platform. Libraries were quantified with Biotium Accuclear Ultra high sensitivity dsDNA Quantitative kit using Mosquito LV liquid handling platform, Bravo WS and BMG FLUOstar Omega plate reader. Libraries were pooled in equimolar amounts on a Beckman BioMek NX-8 liquid handling platform. Libraries were normalised to 2.8 nM ready for cluster generation on a c-BOT and loading on requested Illumina sequencing platform.

#### SNP analysis

SNP analysis was performed using Breseq (76–78). This is a pipeline that performs all the functions including mapping reads using Bowtie2 (79) and calling SNPs and other genomic variations.

The SL1344 parental strain used in this study has two previously described SNPs, in *manX* (E95V) and *menC* (L148L), compared to the reference genome (80) (Fig. S2). While no additional SNPs were identified in WT or GX, the OX strain contained additional SNPs in the genes *ybiH* (V173L), *SL1344_3357* (C260C), and in the intergenic region between *SL1344_3765* and *emrD* (Fig. S2). None of the genes with SNPs were found to interact with either Fis or Dps by Search Tool for the Retrieval of Interacting Genes/Proteins (STRING) analysis (81).

### Revealing transcriptomic changes with RNA-seq

#### RNA extraction

At the EE or TS phases of growth, cells were harvested as described for RT-qPCR. The bacterial pellet was dissolved in TE pH 8.0 (100 mM Tris-HCl pH 8.0, 10 mM EDTA pH 8.0) containing 0.5 mg/ml lysozyme. Lysis was carried out with 1% SDS and 0.1 mg/ml proteinase K while incubating at 40°C for 20 min. RNA was isolated by the addition of 0.3 M sodium acetate (pH 6.5) and 1 volume of phenol (pH 4.3):chloroform (1:1), followed by centrifugation (16000 x g, 4 °C, 10 min) in a phase lock tube (Quantbio, VWR) to separate the aqueous and organic phases. Following this, 1 volume of chloroform was added to the aqueous layer and centrifugation (16000 × g, 4 °C, 10 min) was repeated in the same phase lock tube. RNA was precipitated by the addition of 5 volumes of 100% ethanol and incubation at -20 °C for 1 h. RNA was pelleted by centrifugation (16000 x g, 4 °C, 10 min), washed once with 70% ethanol, and resuspended in 50 µl DEPC treated water. RNA was diluted to 500 ng/µl, denatured at 65°C for 5 min and treated with 10 U RNase-free DNase I (ThermoFisher Scientific, Waltham, USA) in DNase 1 buffer at 37 °C for 40 min. DNase treated RNA was cleaned up using the phenol chloroform method again. RNA integrity was assessed as described previously under the RT-qPCR head.

#### Library preparation

Strand specific library preparation and sequencing of the DNase treated RNA was performed by Vertis Biotechnologie AG (Freising-Weihenstefan, Germany). Briefly, samples were analysed by capillary electrophoresis (Bioanalyzer, Agilent), and ribosomal RNA (rRNA) was depleted using the bacterial Ribo-Zero rRNA Removal Kit (Illumina). Ribodepleted RNA samples were fragmented using ultrasound (4 x 30 second pulses at 4 °C).

Oligonucleotide sequencing adapters were ligated to the 3’ end of each specific strand of the RNA molecules. First-strand cDNA synthesis was performed using M-MLV reverse transcriptase and the 3’ adapter as primer. The first-strand cDNA was purified and the 5’ Illumina TruSeq sequencing adapter was ligated to the 3’ end of the antisense cDNA in a strand specific manner. The resulting cDNA was PCR-amplified to about 10-20 ng/µl using a high fidelity DNA polymerase. The cDNA was purified using the Agencourt AMPure XP kit (Beckman Coulter Genomics) and was analyzed by capillary electrophoresis (Bioanalyzer, Agilent). For Illumina NextSeq sequencing, the samples were pooled in approximately equimolar amounts. The cDNA pool was size fractionated in the size range of 200 - 550 bp using a differential clean-up with the Agencourt AMPure kit. An aliquot of the size fractionated pool was analyzed by capillary electrophoresis. Samples were PCR amplified for Truseq according to the instructions of Illumina. For the WT and GX strains, the cDNA pool was paired end sequenced on an Illumina NextSeq 500 system using 2 x 75 bp read length. For the OX strain, the cDNA pool was single end sequenced on an Illumina NextSeq 500 system using 1 x 75 bp read length.

#### Sequencing alignment

Raw read sequence qualities were assessed using FastQC (http://www.bioinformatics.babraham.ac.uk/projects/fastqc/). BWA (82) was used to align reads to the *Salmonella enterica* subsp. *enterica* serovar Typhimurium str. SL1344 chromosome and plasmids reference sequences (chromosome: NC_016810.1, pCol1B9_SL1344: NC_017718.1, pRSF1010_SL1344: NC_017719.1, pSLT_SL1344: NC_017720.1). Aligned reads were sorted by the reference sequence coordinate and low mapping quality reads (mapping quality < 30) were removed using Samtools (83). Samtools was also used to summarize mapping statistics. More than 90% of the reads were mapped and of the mapped reads, >98% were unique. Sequence coverages were calculated using Deeptools (84) and are available as BigWig (.bw) files that can be viewed using IGB (85).

#### Differential expression analysis

Counts of the number of reads mapping to genomic features (genes or sRNA) were obtained using the R package Rsubread (86). The R package EdgeR (87, 88) was used to identify differentially expressed genes. P values were adjusted using the Benjamini Hochberg method (False Discovery Rate, FDR) and a cutoff of 0.05 was used to call differentially expressed genes. These are available in an Excel file on GEO.

### Revealing Fis binding changes with ChIP-seq

#### ChIP (Chromatin Immunoprecipitation)

For all ChIP experiments *fis*::*8xmyc* epitope tagged variants of WT, GX and OX were used (89). Overnight cultures were diluted to a starting A_600_ of 0.003 in 25 ml LB broth and grown to an A_600_ of 0.4. Cells were harvested by centrifugation at 3,220 × g for 10 min at room temperature and resuspended in 50 ml PBS. Formaldehyde was added dropwise to a final concentration of 1% (v/v) while continuously stirring the cells. After 10 min of crosslinking the reaction was quenched with cold glycine (2M) added dropwise to a final concentration of 0.125 M. Cells were stirred further for 5 min and then harvested by centrifugation at 3220 x g for 10 min at 4 °C. The pellet was suspended in 600 µl Lysis buffer (50 mM Tris-HCl pH 8.1, 10 mM EDTA pH 8.0, 1% (w/v) SDS, Roche protease inhibitor cocktail), incubated for 10 min on ice, prior to the addition of 1.2 ml IP Dilution Buffer (20 mM Tris-HCl pH 8.1, 150 mM NaCl, 2 mM EDTA pH 8.0, 1% (v/v) Triton X-100, 0.01% (w/v) SDS, Roche protease inhibitor cocktail) and transferred to a sonication tube. Chromatin was sheared to an average length of 500 bp by sonicating for 30 seconds with an amplitude of 10 µm, twelve times, with a 1 min incubation on ice between bursts using a Soniprep 150. Sheared chromatin was analysed by agarose gel electrophoresis and stored at -80 °C. To pre-clear chromatin for input samples, 50 µl normal rabbit IgG (Millipore, Cork, Ireland) was added and samples were incubated for 1 h on a blood tube rotator SB1 at 4 °C. Antibody was removed by the addition of 100 µl homogeneous Protein G-Agarose (Roche) and a further incubation for 3 h. The Protein G-Agarose was removed by centrifugation at 3,220 × g for 2 min and 200 µl was used as input. For all ChIP experiments a mock IP and an experimental IP were set up at the same time using 1,350 µl pre-cleared chromatin and 10 µg of normal mouse IgG (Millipore) and monoclonal mouse anti-c-Myc respectively. Samples were incubated at 4 °C for 16 h on a blood tube rotator SB1. 100 µl homogeneous Protein G-Agarose was added to each and the samples were incubated for a further 3 h. Protein G-Agarose beads were pelleted by centrifugation at 5,200 x g, washed 4 times with high salt IP wash buffer (50 mM HEPES pH 7.9, 500 mM NaCl, 1 mM EDTA, 0.1% (w/v) SDS, 1% (v/v) Triton X-100, 0.1% (w/v) deoxycholate), and twice with TE pH 8.0. Protein-DNA complexes were eluted from the beads twice with IP elution buffer (100 mM NaHCO_3_, 1% (w/v) SDS).

DNA was purified and recovered by performing a standard phenol-chloroform extraction, followed by ethanol precipitation with 5 µg of glycogen (Invitrogen, Cat. no. 10814-010). The DNA pellets of the IP samples were re-suspended (by heating at 37°C) in 50 µl of sterile filtered water for experimental and mock IPs, and 100 µl for the Input DNA samples.

#### Library preparation

ChIP sequencing was performed by Vertis Biotechnologie AG using Illumina NextSeq 500 technology. The DNA samples were first fragmented with ultrasound (2 pulses of 30 sec at 4 °C). After end-repair, TruSeq sequencing adapters were ligated to the DNA fragments. Finally, the DNA was PCR-amplified to about 10-20 ng/µl using a high fidelity DNA polymerase. Aliquots of the PCR amplified libraries were examined by capillary electrophoresis (Bioanalyzer, Agilent). For Illumina NextSeq sequencing, the samples were pooled in approximately equimolar amounts. The DNA pool was eluted in the size range of 250 – 550 bp from a preparative agarose gel. An aliquot of the size fractionated library pool was analyzed by capillary electrophoresis (Bioanalyzer, Agilent). The adapters were designed for TruSeq sequencing according to the instructions of Illumina. The shotgun library pool was sequenced on an Illumina NextSeq 500 system using 75 bp read length.

#### Sequencing alignment

See RNA-seq analysis above.

#### Peak calling

Since we were performing differential binding analysis, we did not exclude duplicate reads as they contain information regarding the binding intensity of the transcription factor being pulled down. MACS2 (90) was used to call peaks against the mock. Each strain (WT, GX and OX) had its own mock sample. We used a custom R script to roughly annotate peaks with the names of neighbouring genes and sRNA. These lists are available as both Excel and .gff files. The sequences of Fis peaks from the WT were used to search for motifs using unbiased MEME (91). This Fis motif was then used to search for all possible motifs in the genome, bound or not in the ChIP, using FIMO (92). This list is also available as a .gff file.

#### Differential binding analysis

Differential binding analysis was performed in R (93) using the DiffBind package (94). To correct for differences in gene dosage gradient between strains, mocks derived from each strain were used as control. The output of the differential binding analysis is available (both with and without applying the FDR cutoff) as Excel files.

### *Salmonella* Pathogenicity Island expression analyses using P*gfp*^+^ reporter fusions

SPI-1 (P_*prgH*_ -*gfp*^+^) and SPI-2 (P_*ssaG*_ -*gfp*^+^) promoter fusions were integrated into the SL1344 by transduction by bacteriophage P22, and selected for on LB agar plates supplemented with 25 µg/ml chloramphenicol (95). Overnight cultures of SPI-1 and SPI-2 reporter fusion strains were diluted 1:100 in LB. These were transferred to the wells of a black 96 well plate with a flat and transparent bottom (Corning, Fisher Scientific) in 4-6 technical replicates. The plate was incubated at 37 °C for 24 h with 300 rpm (5mm) shaking in the Synergy H1 microplate reader (Biotek, Vermont, USA). A_600_ and fluorescence (Excitation: 485 nm, Emission: 528 nm) were measured every 20 min. Data were obtained from multiple biological replicates.

### Epithelial cells invasion assay

The gentamicin protection assay as described in (96) was used to determine *Salmonella* invasion. HeLa cells (ATCC) were maintained by passaging in Complete Growth Medium (CGM: Minimum Essential Medium Eagle supplemented with 10% v/v Fetal Bovine Serum, 2 mM L-Glutamine, 1 mM Sodium pyruvate) every 3-4 days (37 °C, 5% CO_2_). HeLa cells were seeded 20-24 h prior to invasion assay in the wells of a 24-well tissue culture plate at a density of ∼5 × 10^4^ cells in 1 ml CGM.

Overnight cultures (16 h growth at 37 °C, 200 rpm) of the *Salmonella* WT, GX, OX, Δ*fis* and Δ*dps* strains were diluted 1:33 in 10 ml LB in a 125 ml flask and grown for 3.5 h (early time point) or 6 h (late time point) at 37 °C, 200 rpm. Cells were pelleted by centrifugation at 8000 × g for 2 min. 900 µl of the supernatant was discarded without disturbing the pellet and then resuspended in 900 µl Hank’s buffered saline solution (HBSS -/-). Cells were diluted 1:10 in HBSS -/- and then diluted in CGM to form the infection mix that was used to infect the HeLa cells at a multiplicity of infection (MOI) of 50 with two replicates per strain. Not more than three strains were tested at a time, with the WT always being tested as the reference strain. The infection was allowed to proceed for 10 min at 37 °C, 5% CO_2_, following which the infection mix was discarded by rapid aspiration. Cells were washed rapidly with HBSS supplemented with calcium chloride and magnesium chloride (+/+) twice to get rid of extracellular bacteria and covered with 1 ml CGM. These were incubated at 37 °C, 5% CO_2_ for 20 min and then again washed rapidly with HBSS +/+ twice and covered with 1 ml CGM supplemented with 50 µg/ml gentamicin to kill any remaining extracellular bacteria. After 1 h incubation at 37 °C, 5% CO_2_ cells were washed rapidly with phosphate buffered saline (PBS) twice and lysed with 1 ml 2% (w/v) sodium deoxycholate in PBS.

Viable counts of the subculture, infection mix and lysed cells were obtained. Percentage (%) invasion was obtained by calculating the % of bacteria in the infection mix that could invade the cells relative to the WT (counts of intracellular bacteria were calculated relative to the counts of the infection mix, to control for variation in infection load, and then relative to the wild type to control for day to day variation). The one-sample t-test was used to assess significance relative to the wild type (µ = 100%, 100% being the % invasion of the WT) and the two-sample Welch t-test was used to assess significance of pairwise comparisons with α = 0.05. The P-values from the two-sample tests were corrected for multiple testing using the BH method (FDR).

### Data availability

RNA-seq and ChIP-seq data can be found on the Gene Expression Omnibus (accession number: GSE152228). WGS data can be found on the Sequence Read Archive (accession number: PRJNA638833). WGS data for the two strains, OX and 2xdO, can be found on the European Nucleotide Archive (accession number: ERS2515905 and ERS4653304, respectively). Scripts used in the analysis can be found on GitLab (<u>Project ID: 10206830</u>).

## Supporting information

Bogue et al. Fis Dps 2020_supplementary_files

## ACKNOWLEDGEMENTS

We thank Prof. Dr. Regine Hengge (Humboldt-Universität zu Berlin) for providing the Rabbit-α-Dps polyclonal serum. We thank Dr. Carsten Kröger (Trinity College Dublin) and Prof. Jay Hinton (University of Liverpool) for providing the P_*prgH*_ and P_*ssaG*_ *gfp*^+^ reporter strains. We thank Dr. Ciaran Finn (NIH, Trinity College Dublin) and Dr. Olivia Steele-Mortimer (NIH) for providing the HeLa cells and helping us with the *Salmonella* invasion assay protocol. We are grateful to Matthew Dorman for assistance with depositing whole genome sequence data to the European Nucleotide Archive. Research in Charles Dorman’s laboratory is supported by Principal Investigator Award 13/1A/1875 from Science Foundation Ireland. Whole genome sequencing at the Wellcome Sanger Institute was supported by grant 206194 from the Wellcome Trust.

