## Supplementary material for "Network rewiring: physiological consequences of reciprocally exchanging the physical locations and growth-phase-dependent expression patterns of the *Salmonella fis* and *dps* genes": Bogue et al. Fis Dps 2020_supplementary_files

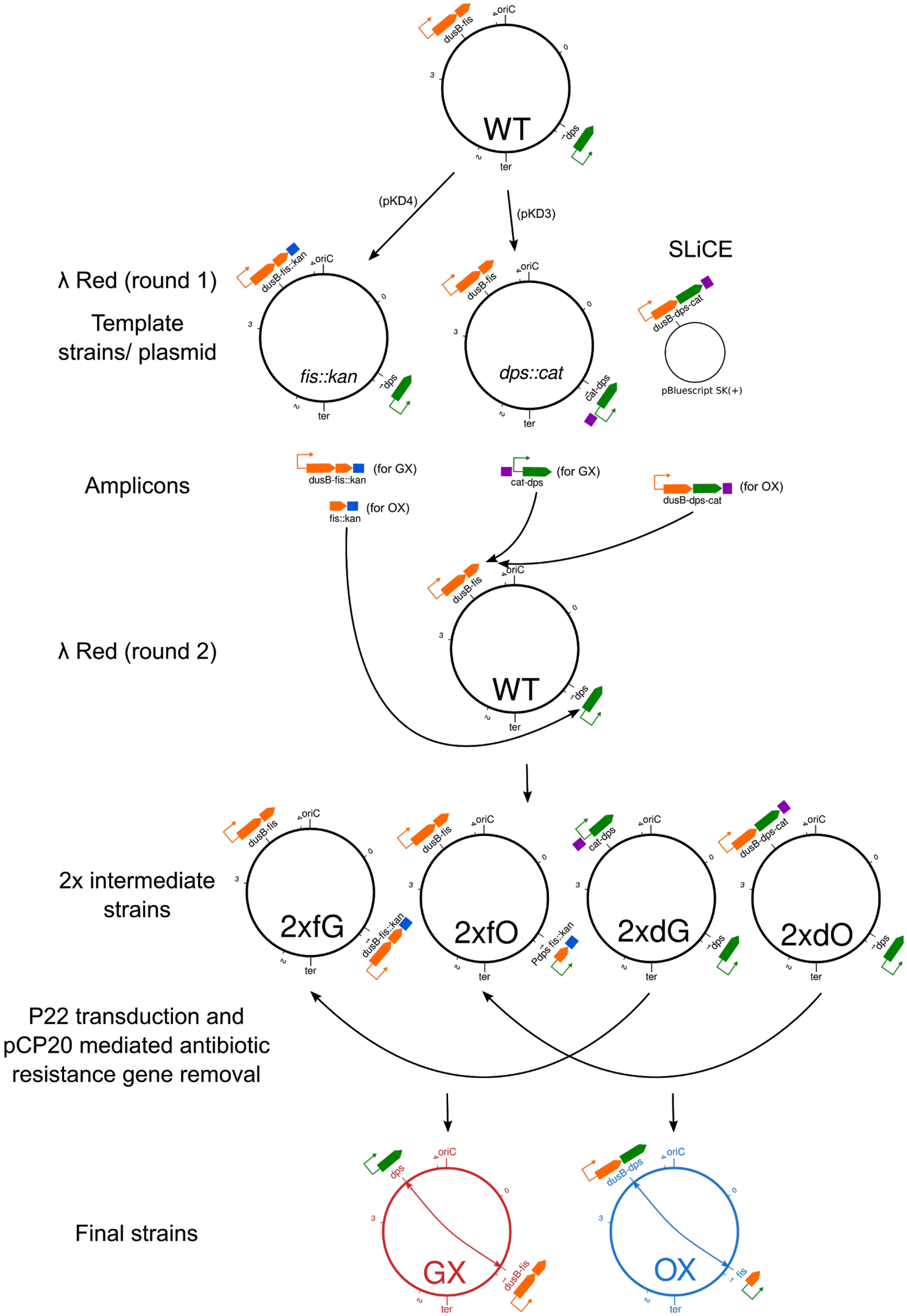


**Figure S1. Schematic representation of the strain construction process.** For description see Materials and Methods.

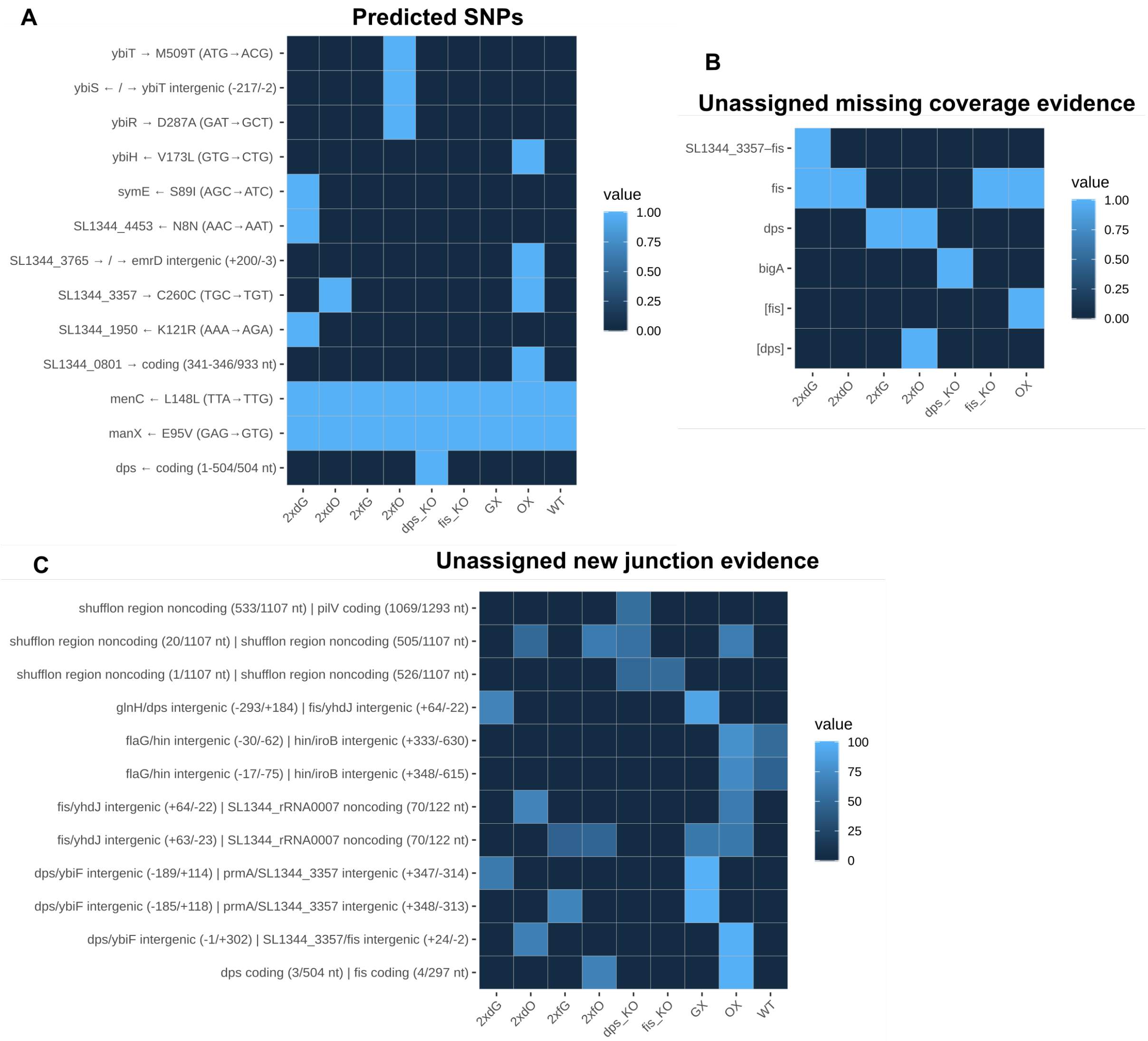


**Figure S2. Summary of SNP analysis on constructed strains.** Heatmaps depicting summarized Breseq analysis results. **(A)** Predicted mutations, **(B)** Unassigned missing coverage evidence and **(C)** Unassigned new junction evidence. In (A) and (B) colour scale indicates presence (1)/ absence (0) of mutation. In (C) colour scale indicates frequency of the new junction among reads mapping to that locus.

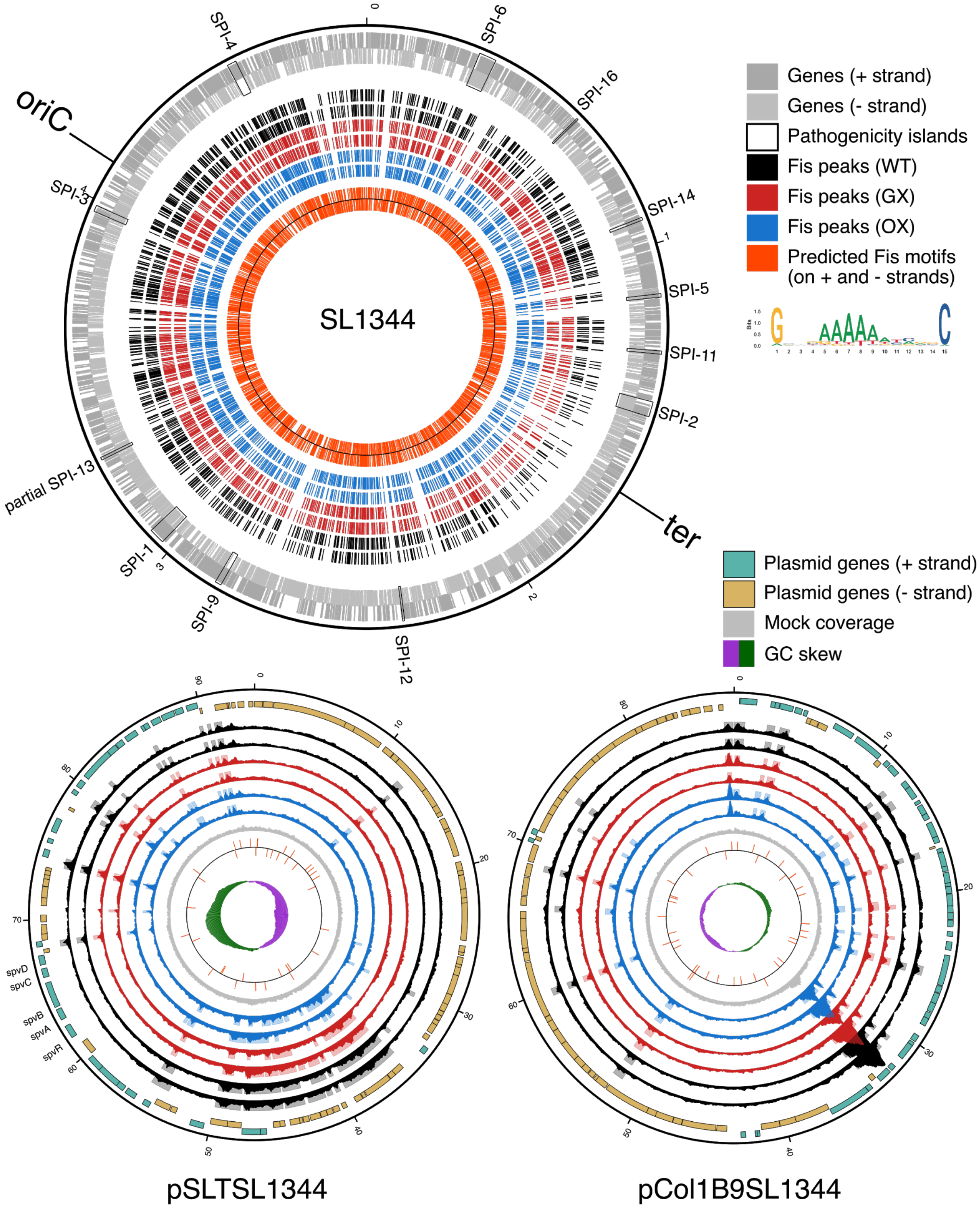


**Figure S3. Fis binding regions on the chromosome and plasmids.** SL1344 chromosome: predicted Fis binding regions are shown for the two biological replicates of WT, GX and OX in early exponential phase. Increased Fis binding around the *ter* in the GX and OX strains results in slightly better peak calling in this region compared to the WT. Plasmids: Coverages from the ChIP-seq experiment (solid traces) and predicted Fis binding regions (shaded boxes) are shown for the two biological replicates of WT, GX and OX in early exponential phase. For comparison, the wild type mock coverage is also shown as a solid grey trace.

Chromosomal positions are in millions of base pairs and plasmid positions are in thousands of base pairs.

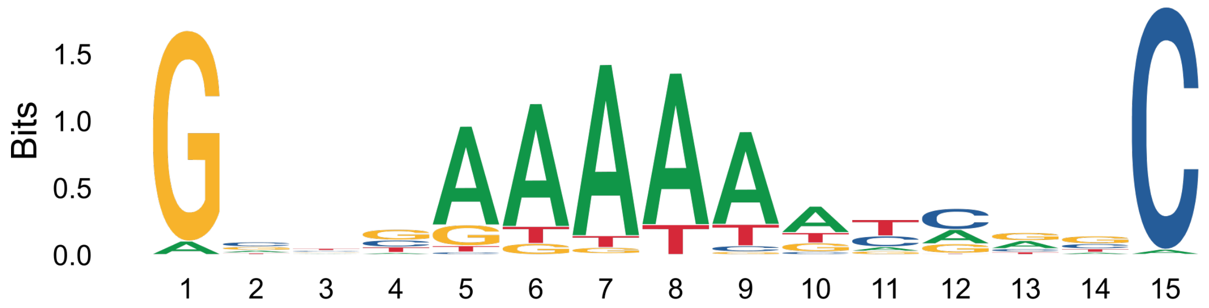


E-value: 3.5e-062
Log Likelihood Ratio: 11054
Site Count: 1461
Information Content: 10.9
Relative Entropy: 10.9
Bayes Threshold: 10.4408
Width: 15

**Figure S4. Fis binding motif obtained from ChIP-seq data.** A 15 nucleotide long sequence logo representing the Fis binding motif discovered using MEME on the set of Fis binding regions in the wild type identified using ChIP-seq. The height of each letter represents the relative frequency of that nucleic acid at that position in the motif. The overall height of the stack of letters at each position represents the degree of conservation at that position in the motif (if the stack is very short, that position is highly variable). Various statistics reported by MEME are presented below the logo. This motif is similar to the *E. coli* Fis binding motif.

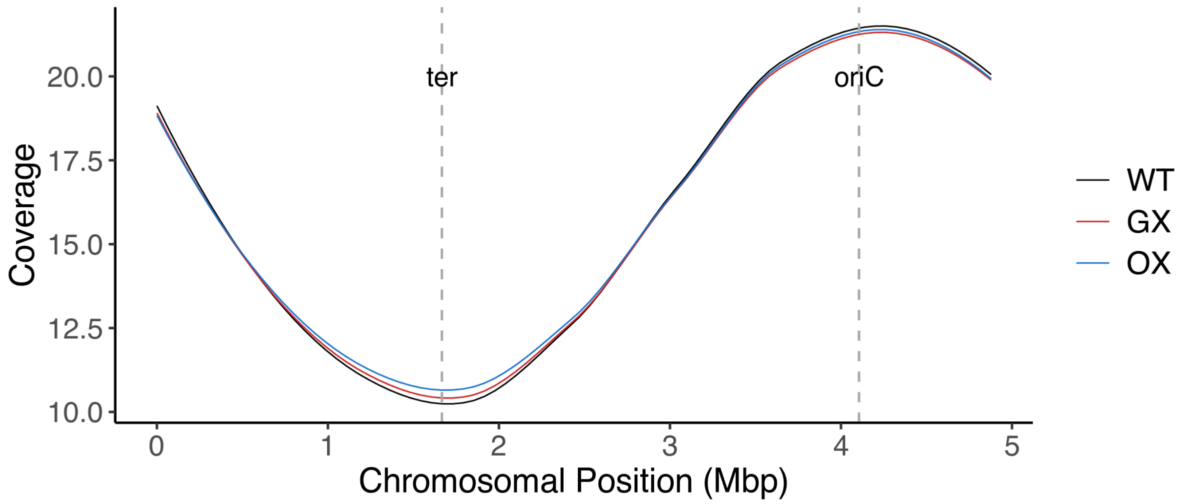


**Figure S5. Gene dosage gradient differences in the GX and OX strains compared to WT visualized using coverage data from the ChIP-seq mocks of these strains.** For clarity, the coverage data at each position is not shown, only the local average calculated using the R loess function along the chromosomal position in millions of base pairs is shown. There are only slight gene dosage gradient differences in the GX and OX strains at the early exponential time point at which chromatin was collected for ChIP-seq. These differences are considered in the calculation of differentially bound regions.

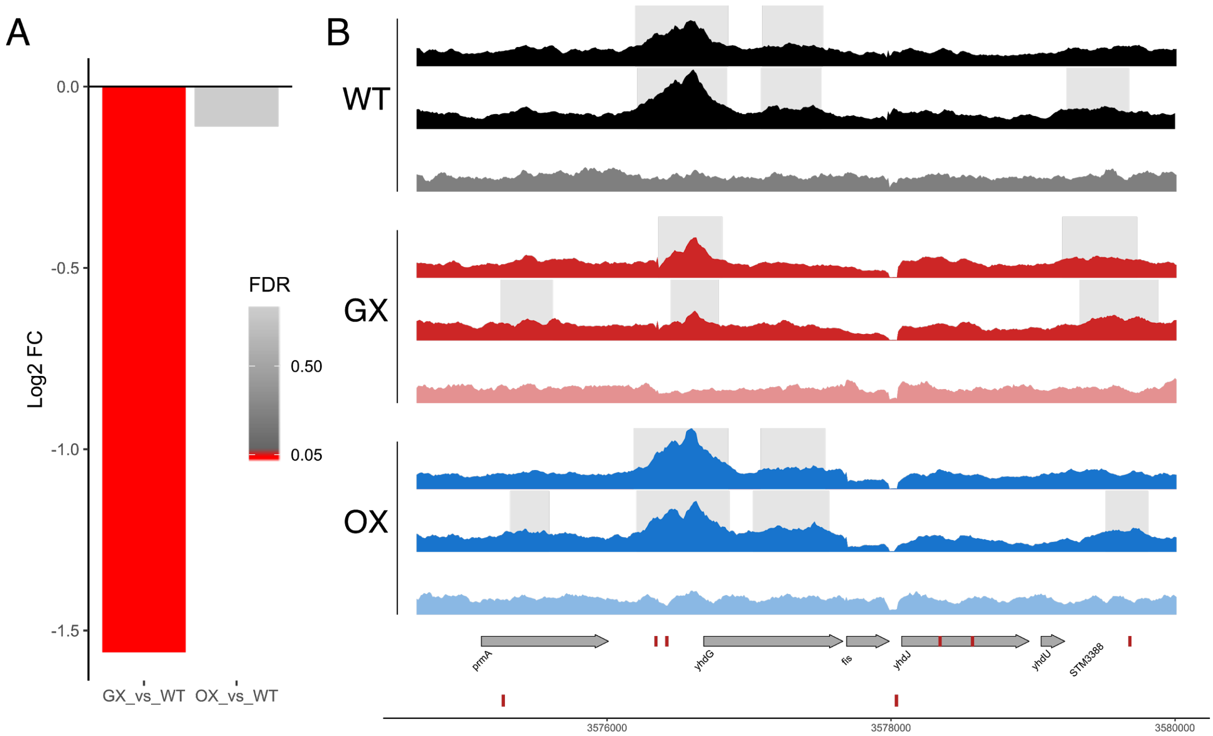


**Figure S6. Fis binding around *fis* gene. (A)** Log$_2$ fold changes of Fis binding upstream of the *dusB-fis* transcription unit in GX and OX strains relative to the WT. **(B)** Fis ChIP-seq coverages of the WT, GX and OX strains (two biological replicates and mock) of the region around the *fis* gene. Grey shaded regions show peaks called in each sample relative to the mock. Genes (grey arrows) and Fis binding motifs (red blocks) are shown on the + and - reference strands. The *fis* gene is a part of the *dusB-fis* transcription unit. The promoter of this transcription unit contains two Fis binding motifs, as predicted by FIMO from the MEME suite, and in the ChIP-seq data shows substantial Fis binding. ChIP-seq data was analysed using the wildtype reference sequence and thus the coverage here is plotted relative to wildtype gene locations, even though these locations have been altered in the GX and OX strains. In the GX strain, the entire region including *dusB*, *fis* and 315 bp of its upstream promoter region has been relocated to the *dps* locus. In the OX strain only the *fis* ORF has been relocated to the *dps* locus.

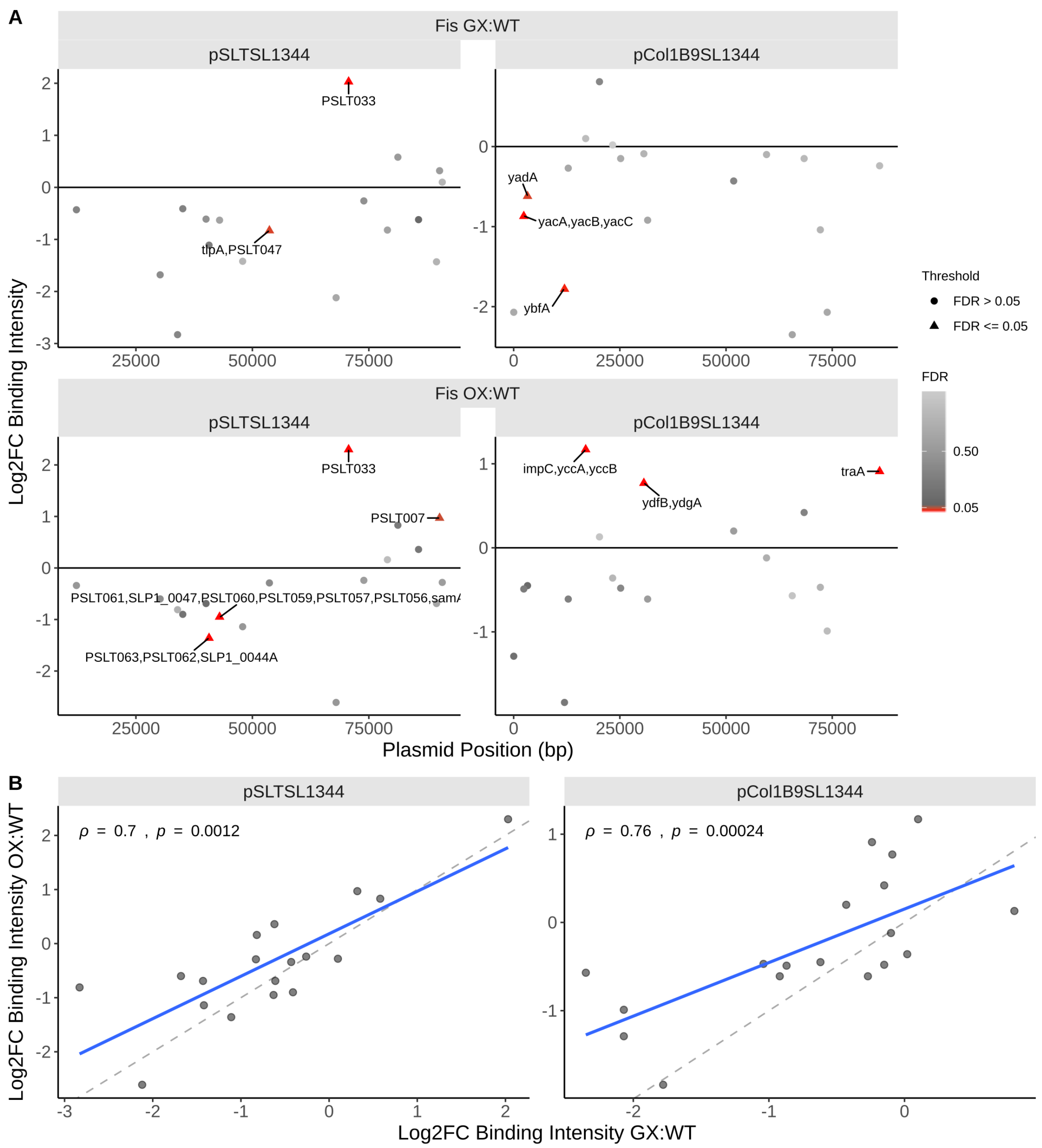


**Figure S7. Fis binding changes on the *Salmonella* megaplasmids assayed by ChIP-seq during EE phase. (A)** Peak intensity changes along the plasmid position shown in base pairs. In calculating these intensity changes, the appropriate mocks (WT mock for WT peaks, GX mock for GX peaks, OX mock for OX peaks) were used as control. Significantly differentially bound peaks are roughly annotated with genes that are located in their vicinity. **(B)** Spearman's correlation of peak intensity changes in the GX with the OX strain. Dashed line has slope of 1 and indicates the position where the fold changes in Fis binding intensity in GX would be equal to those in OX. Blue solid line is the regression linear fit to the data.

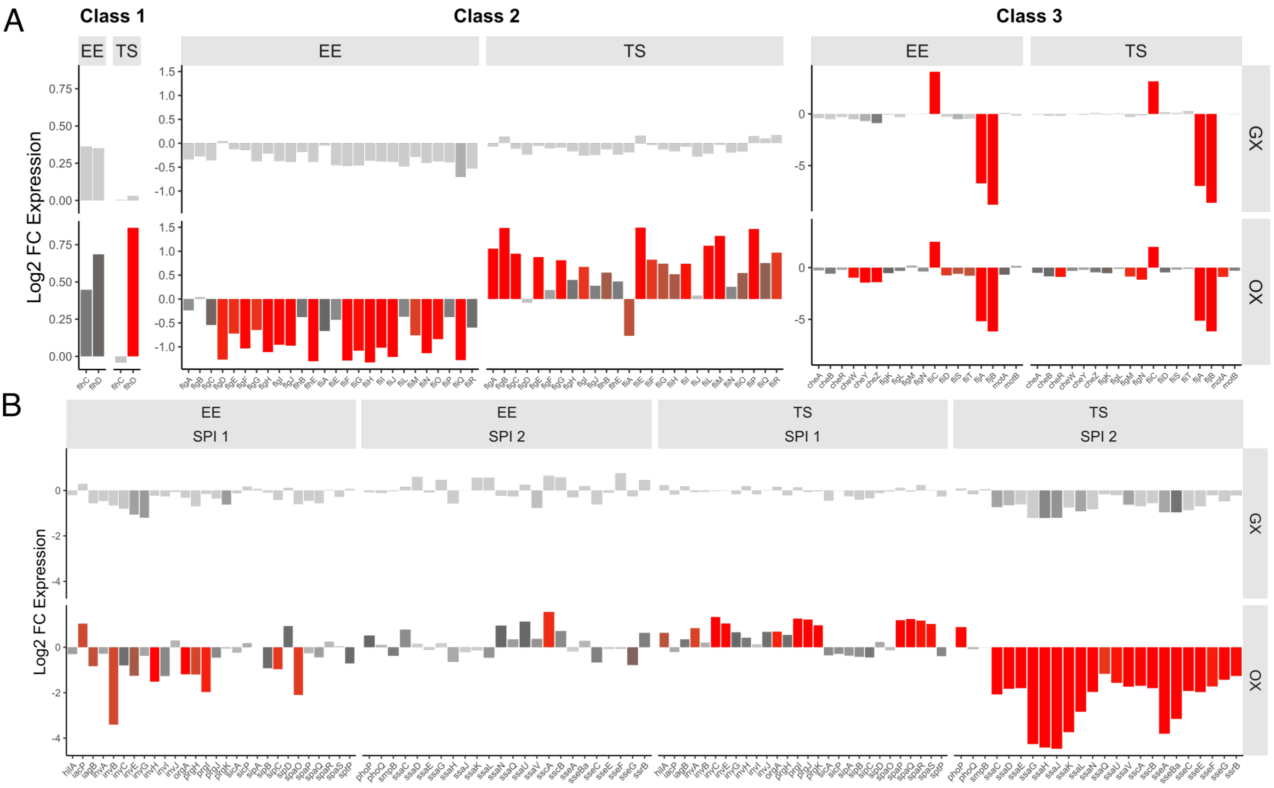


**Figure S8. Fold changes of some gene sets in the GX and OX strains. (A)** Motility genes and **(B)** SPI-1 and 2 genes. Expression changes in early exponential (EE) and transition to stationary (TS) phases are shown. None of these gene sets are significantly differentially expressed in the GX strain. Note that some of the Class 3 motility genes (*fliC, fljA* and *fljB*) were differentially expressed due to phase switching in the GX and OX strains during their construction. The orientation of these genes was different in the constructed strains with respect to the wildtype against which gene expression changes were calculated, thus resulting in phase-variation related gene expression changes. These changes are unlikely to be related to our study. A general pattern of reversal in gene expression between the early exponential and transition to stationary phases was seen in the OX strain (this could be related to Fis levels being higher in the stationary phase in this strain). Colour bar indicates FDR and bars representing genes with FDR < 0.05 are red.

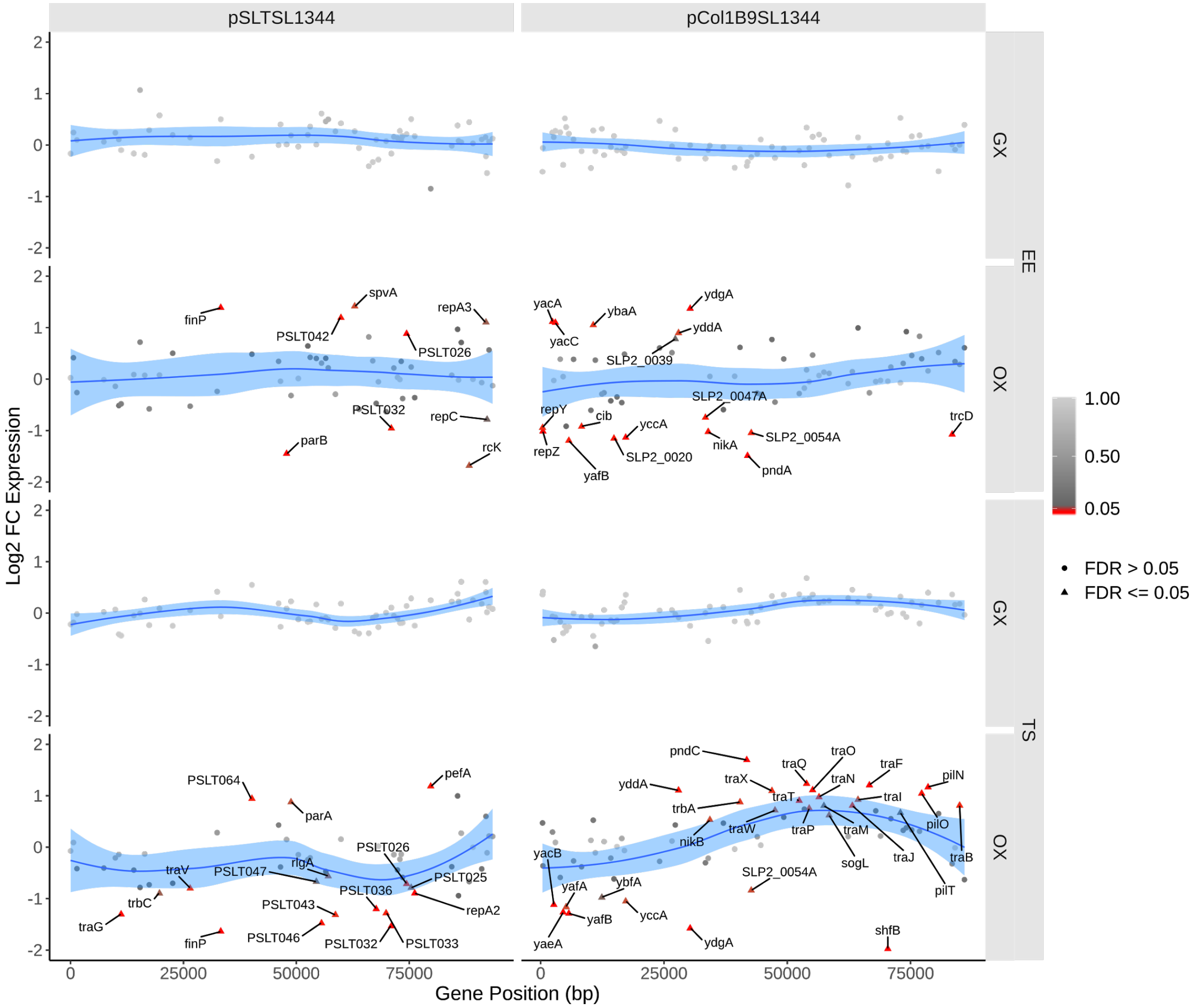


**Figure S9. Transcriptomic changes on the *Salmonella* megaplasmids assayed by RNA-seq.** $Log_2$ fold changes in expression of genes along their position on the plasmid in base pairs. The blue curve shows the local regression peak intensity change calculated using the R loess function. Significant gene expression changes (FDR < 0.05) are shown as red triangles.

**Table S1 Strains used in this study.**

| Strain | Genotype | Description |
|---|---|---|
| SL1344 | *rpsL* *hisG46* *manX*E95V *menC*L148L | (1, 2) |
| SL1344 *fis*::*kan* | *fis*::*kan* | Kanamycin resistance cassette inserted 63 bp downstream of fis ORF |
| SL1344 *dps*::*cat* | $P_{dps}$::*cat* | Chloramphenicol resistance cassette inserted 149 bp upstream of dps ORF |
| SL1344 2xfG | Δ$P_{dps}$*dps*::$P_{fis}$*dusBfis* | *dps* ORF and upstream/downstream regulatory regions replaced with *dusBfis* operon and regulatory regions |
| SL1344 2xdG | Δ$P_{fis}$*dusBfis*::$P_{dps}$*dps* | *dusB* and *fis* ORFs and upstream/downstream regulatory regions replaced with *dps* ORF and regulatory regions |
| SL1344 2xfO | Δ*dps*::*fis* | *dps* ORF replaced with *fis* ORF |
| SL1344 2xdO | Δ*fis*::*dps* | *fis* ORF replaced with *dps* |
| SL1344 Δ*fis* | Δ*fis*::*cat* | (3) |
| SL1344 Δ*dps* | Δ*dps* | *dps* ORF deletion |
| SL1344 GX | Δ$P_{dps}$*dps*::$P_{fis}$*dusBfis* Δ$P_{fis}$*dusBfis*::$P_{dps}$*dps* | Exchange of *dusBfis* and *dps* and their *cis* regulatory regions |
| SL1344 OX | Δ*dps*::*fis* Δ*fis*::*dps* | Exchange of *fis* and *dps* ORFs |

| | | |
|---|---|---|
| SL1344 *fis*::*8xmyc* | *fis*::*8xmyc* | Deletion of the *fis* stop codon and insertion of 8 copies of the Myc epitope tag separated by a 21 bp linker. |
| SL1344 GX *fis*::*8xmyc* | GX *fis*::*8xmyc* | Deletion of the *fis* stop codon and insertion of 8 copies of the Myc epitope tag separated by a 21 bp linker. |
| SL1344 OX *fis*::*8xmyc* | OX *fis*::*8xmyc* | Deletion of the *fis* stop codon and insertion of 8 copies of the Myc epitope tag separated by a 21 bp linker. |
| SL1344 P*prgH* | P*prgH*::*gfp+*::*cat* | GFP+ fused to SPI-1 promoter, P*prgH* (4) |
| SL1344 GX P*prgH* | P*prgH*::*gfp+*::*cat* | GFP+ fused to SPI-1 promoter, P*prgH* |
| SL1344 OX P*prgH* | P*prgH*::*gfp+*::*cat* | GFP+ fused to SPI-1 promoter, P*prgH* |
| SL1344 Δ*fis* P*prgH* | P*prgH*::*gfp+*::*cat* | GFP+ fused to SPI-1 promoter, P*prgH* |
| SL1344 Δ*dps* P*prgH* | P*prgH*::*gfp+*::*cat* | GFP+ fused to SPI-1 promoter, P*prgH* |
| SL1344 P*ssaG* | P*ssaG*::*gfp+*::*cat* | GFP+ fused to SPI-2 promoter, P*ssaG* (4) |
| SL1344 GX P*ssaG* | P*ssaG*::*gfp+*::*cat* | GFP+ fused to SPI-2 promoter, P*ssaG* |
| SL1344 OX P*ssaG* | P*ssaG*::*gfp+*::*cat* | GFP+ fused to SPI-2 promoter, P*ssaG* |
| SL1344 Δ*fis* P*ssaG* | P*ssaG*::*gfp+*::*cat* | GFP+ fused to SPI-2 promoter, P*ssaG* |
| SL1344 Δ*dps* P*ssaG* | P*ssaG*::*gfp+*::*cat* | GFP+ fused to SPI-2 promoter, P*ssaG* |
| DH5α | *fhuA2 lacΔU169 phoA glnV44 Φ80' lacZΔM15* | (5) |

| | | |
|---|---|---|
| | *gyrA96 recA1 relA1 endA1 thi-1 hsdR17* | |
| DH5α pBLUESCRIPT-*dusB-dps-cat* | pBLUESCRIPTSK(+)::*dusB*::*dps*::*cat* | *dusB-dps* operon with *cat* antibiotic resistance gene |

**Table S2 Oligonucleotides used in this study.**

| Name | Sequence 5'-3' |
|---|---|
| fisVal-F | GCTGTCCGGGTTGTTCTG |
| fisVal-R | ACCAAATTCCATGTGATGCGT |
| dpsVal-F | CAGTATGCCGCACCGTTT |
| dpsVal-R | GCGCTATTACTTCGTCATTTTTTGT |
| fiskanInsert-F | TAAAAAGGCGCTACTCGGCATGGGGAAGCGCC TTTTTTATGTGCCACCTGCATCGATGGC |
| fiskanInsert-R | TTCACATTCCGCTTTCATGACCAAATTCCATGT GATGCGTCATATGAATATCCTCCTTAG |
| dpscmInsert-F | GACGAGTTTGCTGTTTGGTGGGTAATAATTCTC ATTTTAAGTGTAGGCTGGAGCTGCTTC |
| dpscmInsert-R | TACTTCGTCATTTTTTGTCATATTTTTTCTCATTT TTTACCATATGAATATCCTCCTTA |
| fisReg-F | GTCATTTTTTGTCATATTTTTTCTCATTTTTTACA TTAAAGTGATCTTGTCCTGAAAC |
| fisReg-R | CAAATAATTCACTTTTGTCGGAGGGGAGTACAA GACGTGTCATATGAATATCCTCCTTAG |
| fisORF-F | TTAATTACCTGGGACACAAACATCAAGAGGATA TGAGATTATGTTCGAACAACGCGTA |
| dpsReg-F | TTCACATTCCGCTTTCATGACCAAATTCCATGT GATGCGTCAAGACGTGTGCACTATT |
| dpsReg-R | CTTCCGACTGGCAATGGAGAAAAATCACGCGC AGCGGGAACATATGAATATCCTCCTTAG |
| pBLUE_BB_F | CCAGCTTTTGTTCCCTTTAG |
| pBLUE_BB_R | CCCAATTCGCCCTATAGTG |
| pBLUE_dusB_F | GTGAATTGTAATACGACTCACTATAGGGCGAAT TGGGCGATTCATTGATCTACAACA |
| pBLUE_dusB_R | GATTAGACGCTTTTGTTTTTACCAGTTTAGCGG TACTCATAGTTCTGTCAGCTCTTTATTTC |
| pBLUE_dps_F | GAAAATTTTGCGTAAACAGAAATAAAGAGCTGA CAGAACTATGAGTACCGCTAAACTGGT |
| pBLUE_dps_R | TAGGAACTTCGGAATAGGAACTAAGGAGGATA TTCATATGCAAGACGTGTGCACTATTTA |
| pBLUE_cat_F | TGGCACAGGGGTTTTGCACTTAAATAGTGCAC ACGTCTTGCATATGAATATCCTCCTTAG |

| | |
|---|---|
| | AAGCTCGAAATTAACCCTCACTAAAGGGAACAA |
| pBLUE_cat_R | AAGCTGGGTGCCACCTGCATCGATGGC |
| | GGTACGCTGCGTAAAAAATTAAAAAAATACGGC |
| fis8mycins_F | ATGAACGTCGGATCCAGTCTTCGTGAT |
| | GAGTAGCGCCTTTTTAAACAAGCAGTTAGCTAA |
| fis8mycins_R | TCGAAAAATTCCGGGGATCCGTCGACC |
| pBLUE_chq_F | GACGTTGTAAAACGACGGCC |
| pBLUE_chq_R | CGGCTCGTATGTTGTGTGG |
| | ATTTTAATGAATTATAAATTTATTTTGCTGTTTTT |
| SLiCE_dpsORF_F | AAGCACGATTCATTGATCTACAACA |
| | TTCACATTCCGCTTTCATGACCAAATTCCATGT |
| SLiCE_dpsORF_R | GATGCGTGTGCCACCTGCATCGATGGC |

**Table S3 Oligonucleotides used for qPCR.**

| Name | Sequence 5’-3’ |
|---|---|
| RT_fis_F | TGACGTACTGACCGTTTCTACCGT |
| RT_fis_R | ACGTCCTGACCATTCAGTTGAGCA |
| RT_dps_F | CACTGACCGATCATCTGGATAC |
| RT_dps_R | CGGATAGCTTTTCAGTGGAGTT |
| RT_hemX_F | CGCCTGACGGTATGTTTCTT |
| RT_hemX_R | CCCAACCAGGACGTCTATTTAC |
| RT_gidA_F | CAGATTCGGCTGATTCTCCA |
| RT_gidA_R | TCAGGCAGGTATTCAGTTTAGG |
| RT_STM1554_F | TTTAGACGCCCGCACTTC |
| RT_STM1554_R | TAGCTCTCCCGAGTTAGATAATCA |
